# Rational design of ^19^F NMR labelling sites to probe protein structure and interactions

**DOI:** 10.1101/2024.12.11.627779

**Authors:** Julian O. Streit, Sammy H.S. Chan, Saifu Daya, John Christodoulou

## Abstract

Proteins are investigated in increasingly more complex biological systems, where ^19^F NMR is proving highly advantageous due to its high gyromagnetic ratio and background-free spectra. Its application has, however, been hindered by limited chemical shift dispersions and an incomprehensive relationship between chemical shifts and protein structure. We exploit the sensitivity of ^19^F chemical shifts to ring currents by designing labels with direct contact to a native or engineered aromatic ring. Fifty protein variants predicted by AlphaFold and molecular dynamics simulations show 80-90% success rates and direct correlations of their experimental chemical shifts with the magnitude of the engineered ring current. Our method consequently improves the chemical shift dispersion and through simple 1D experiments enables structural analyses of alternative conformational states, including ribosome-bound folding intermediates, and in-cell measurements of thermodynamics and protein-protein interactions. Our strategy thus provides a simple and sensitive tool to extract residue contact restraints from chemical shifts for previously intractable systems.

## Introduction

NMR spectroscopy is a powerful experimental technique to probe biomolecular structure, interactions, and dynamics across different timescales. As a deeper understanding of protein structure within larger complexes and its physiological environment is increasingly being sought, ^19^F NMR has re-emerged as a valuable tool in contemporary structural biology, with continuing methodological advancements permitting the study of systems that are inaccessible by other means^1–5^.

The fluorine nucleus is highly attractive for biomolecular NMR, as its 100% natural abundance and high gyromagnetic ratio (94% of ^1^H) ensures high (signal-to-noise) sensitivity, while it is relatively inexpensive relative to other labelling schemes. Residue-type or site-specific labelling by biosynthetic means or covalent modification results in background-free labelling and reduced spectral complexity that permits the use of simple 1D pulse-acquire experiments. Thus, lower sample concentrations are required, with greater spectral observability of larger molecules and challenging systems, including membrane proteins^2,3,5–7^, nascent polypeptides in complex with their ∼2.5-MDa parent ribosome^8,9^, and within living bacterial^10,11^ and mammalian cells^12,13^. Fluorine NMR is also commonly used in drug screening^14^.

The potential of ^19^F NMR to further advance our understanding of structural biology is, however, constrained by two main limitations. Chemical shift dispersion is often limited in complex biological systems, precluding the ability to resolve multiple conformational states or labelling sites^2–4,8,15,16^. This is due to line broadening associated with fast spin relaxation of slowly tumbling, large biomolecules, exacerbated in-cell by quinary interactions and decelerated diffusion. The use of trifluoromethyl (CF_3_) groups has become the preferred reporter for proteins because of the high (signal-to-noise) sensitivity from the three-fold degeneracy and reduced chemical shift anisotropy (CSA) from the rotationally mobile CF_3_ group^17^. However, this also results in limited chemical shift dispersions deriving from its reduced sensitivity to local electric fields and Van der Waals interactions compared to monofluorinated tags^3,16^, though new fluorinated amino acids are being developed to improve chemical shift sensitivity^3,16,18^.

A second limitation is that ^19^F NMR is not routinely used to obtain structural restraints. Unlike other NMR nuclei whose chemical shifts can be used to derive structural models^19–22^, the origins of ^19^F protein chemical shifts are not well understood^23–25^ and predictions rely on computationally expensive quantum chemical calculations^26^. Moreover, like many other methods, such as Förster resonance energy transfer (FRET) and double electron-electron resonance (DEER), ^19^F NMR requires protein modification. While ^19^F labels can be considerably less bulky, with analogous fluorinated amino acids available^25^, an important, yet often overlooked, consideration is the choice of labelling site. This therefore provides an opportunity to relate site-specific ^19^F-labels on protein structure with their chemical shifts.

Using the commercially available ^19^F-label 4-trifluoromethyl-L-phenylalanine (tfmF), we present a design method that overcomes these challenges. Motivated by previous reports of aromatic ring currents affecting ^19^F NMR chemical shifts^5,6,27^, we have rationally designed 50 protein variants with fluorine labelling sites near native or engineered aromatic rings. Using AlphaFold^28–31^ and all-atom molecular dynamics (MD) as predictors with 80-90% success, Van der Waals contacts between two sidechains are measured directly as the ^19^F chemical shift from simple 1D NMR spectra. We illustrate potential applications of our approach applied to multiple systems where other experimental methods are unable to provide residue-specific insights.

## Results

### Ring current effects improve ^19^F chemical shift dispersion

To begin exploring the utility of probing different sites by ^19^F NMR, we initially used the model immunoglobulin-like domain FLN5, whose structure and folding *in vitro*^32,33^ and co-translational folding (coTF) on the ribosome^8,9,34,35^ has been well-characterised by NMR spectroscopy, including by ^19^F-labelling^8,36^. We initially chose ^19^F-labelling using the non-canonical amino acid tfmF because it yields NMR spectra with high signal-to-noise for even MDa complexes^8^, is commercially available, and can easily be incorporated into standard bacterial strains^37^. Similar fluorinated phenylalanine analogues have also been incorporated in human cells^12,38^. An efficient in-frame amber suppression protocol (>95% incorporation^8^) was used to incorporate tfmF at genetically specified positions in FLN5 (**Fig. 1a-b**) in *E. coli*. We produced 18 variants of FLN5, each tfmF-labelled at different positions across the whole protein structure (16 solvent-exposed, and 2 within the hydrophobic core, 691tfmF and 747tfmF) and all yielding a single resonance in their ^19^F NMR spectrum. The chemical shifts of the 18 labelling sites ranged across ∼0.4 ppm (**Fig. 1c**), centred at the random coil value (61.82 ppm as observed for three tfmF-labelling sites on an unfolded variant of FLN5, **Fig. 1c**), and thus appear to be significantly narrower in their dispersion compared to the ∼2 ppm chemical shift range of trifluoromethyl-labelled membrane proteins^2–5^, likely due to a more uniform chemical environment in bulk solution compared to membranes.

**Figure 1.**
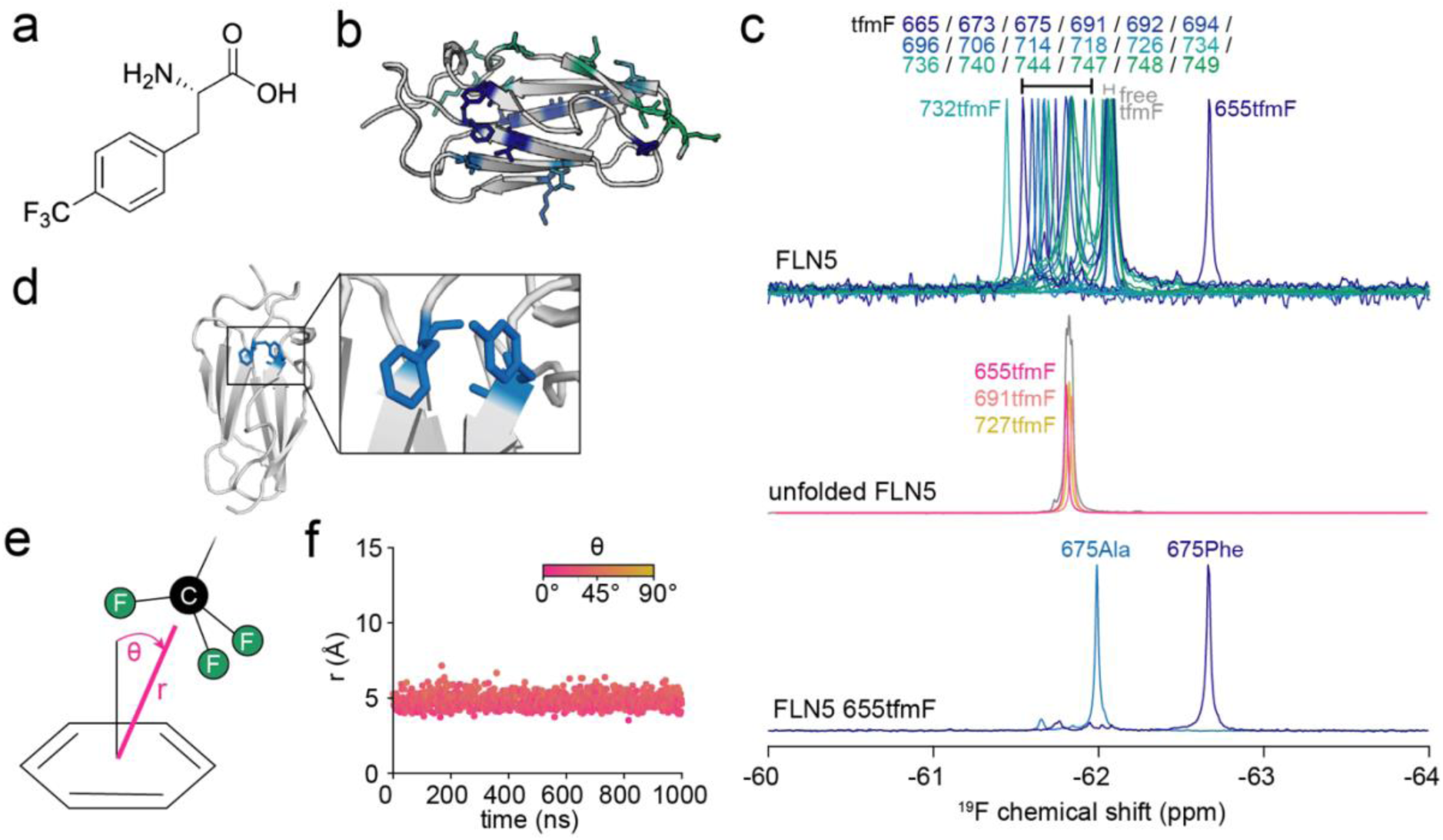
Aromatic ring currents dominate the secondary chemical shift of a fluorinated protein. **a.** Chemical structure of 4-trifluoromethyl-L-phenylalanine (tfmF). **b.** Crystal structure of FLN5 (PDB 1QFH) with residues used for tfmF labelling highlighted (colour scheme corresponds to legend shown in panel c). **c.** ^19^F NMR spectra of folded FLN5 (top) and unfolded FLN5 (A_3_A_3_ mutant^8,34^, middle) labelled at various positions as highlighted in panel **b**. The bottom spectra show wild-type (675Phe) and mutant (Phe675Ala) FLN5 655tfmF. All spectra were recorded at 298 K and 500 MHz. **d.** Close-up view of Tyr655 and Phe675 in the crystal structure of FLN5. **e.** Schematic illustration of a trifluoromethyl (CF_3_) group interacting with an aromatic benzene ring, defined by the distance, *r*, between the CF_3_ and ring centres of mass and the angle, *θ*, between the vector normal to the ring plane and the vector between the CF_3_ and ring centres of mass. **f.** All-atom MD simulation of FLN5 655tfmF showing an interaction between the CF_3_ group of 655tfmF and the aromatic ring of Phe675 quantified using *r* and *θ*.

Two exceptions to this narrow dispersion of chemical shifts are 655tfmF (−62.61 ppm) and 732tfmF (−61.44 ppm). Inspection of the FLN5 crystal structure revealed that both residues are uniquely positioned near residues with a (de)shielding effect. Residue 732 is positioned close to a negatively charged Glu692 on the neighbouring strand, which likely explains the observed deshielding effect (+0.38 ppm) as protein ^19^F chemical shifts have been shown to be sensitive to charge mutations, although the magnitude of electrostatic interactions contributing to shielding remains poorly understood^24^. A larger dispersion was observed for 655tfmF, exhibiting a resonance −0.79 ppm from the random coil. Because residue Tyr655 is in close contact with Phe675 on the adjacent β-strand (**Fig. 1d**), we tested whether a ring current effect induced by Phe675 could cause shielding of the ^19^F nucleus of 655tfmF. Mutation of Phe675 to alanine resulted in only small ^1^H/^15^N amide backbone CSPs, with the protein remaining folded (**Fig. S1a**). However, we observed a significantly less shielded ^19^F chemical shift of −62.0 ppm, within the range of all other labelling sites (**Fig. 1c**). Ring current effects can therefore dominate the secondary chemical shifts of solvent-exposed fluorine labelling sites, and the resulting chemical shifts are thus direct reporters of specific sidechain contacts.

We next assessed site-specific labelling on FLN5 at position 655 using two alternative trifluoromethyl probes. Incorporation of 4-(trifluoromethoxy)-l-phenylalanine (OCF_3_Phe) by amber suppression yields NMR resonances with high signal-to-noise, but with a chemical shift dispersion reduced from that of tfmF by 0.56 ppm (70%, **Fig. S2**). Similarly, post-translational cysteine modification with 2-bromo-N-(4-[trifluoromethyl]phenyl)acetamide (BTFMA) has been shown to be most sensitive to solvent polarity (and likely protein environments) among trifluoromethyl tags^15^, yet for FLN5 Tyr655Cys Cys747Val results in only a < 0.01 ppm secondary chemical shift (**Fig. S3a**). The increased flexibility of both tfmOF and Cys-BTFMA sidechains are likely incompatible with close Phe675 contacts to induce a strong ring current effect (**Fig. S3b-d**).

To quantify the sidechain interactions and thus ring current effects between 655tfmF and 675Phe, we used all-atom molecular dynamics (MD) simulations with two force fields, ff15ipq^39,40^ and C36m^41,42^ (see Methods). Simulations with both force fields corroborate a stable sidechain interaction, with an average distance of ∼5 Å and a perpendicular orientation between the CF_3_ group and the benzyl ring of 675Phe, collectively compatible with a shielded ring current effect as measured experimentally (**Fig. 1e-f, Fig. S1b-c**), and which rationalise the limited secondary chemical shifts observed with tfmOF and BTFMA labelling. This result is independent of the initial sidechain orientation of 655tfmF (**Fig. S1b-c**), suggesting these simulations can be predictive. In the reverse scenario, with 675tfmF and Tyr655, we observe distances that are not compatible with ring current effects (> 7 Å, **Fig. S1b-c**), consistent with an experimentally observed ^19^F chemical shift of −61.67 ppm (deshielded by 0.15 ppm relative to random coil, **Table S3**).

### Rational design of ring current shifts to probe protein structure

We explored whether novel fluorine labelling sites could be designed with structurally interpretable and enhanced secondary chemical shifts using native or engineered ring currents. Given the precise nature of a ring current contact and the labour- and time-consuming effort of an entirely trial-and-error experimental approach, we instead developed a rational design approach exploiting either native aromatics or engineered residues (**Fig. 2a**) and initially applied them to FLN5 and human HRAS. We predicted and quantified sidechain orientations of new tfmF/aromatic-residue labelling pairs using all-atom MD simulations performed in triplicate from different initial sidechain orientations (3 x 1 μs, see Methods, **Table S1, Fig. S4**), and orthogonally from structures predicted by AlphaFold2 (AF2)^28^/ColabFold^29^ and AlphaFold3 (AF3)^31^ (where tfmF is mimicked by Tyr in AF2/AF3). Recent assessments of AlphaFold2 showed that >80% of sidechain conformers predicted with confidence (pLDDT ≥ 70) align with experimental electron density maps^43^. We used the simulated and predicted structures to seek variants possessing short distances between tfmF label and a proximal aromatic residue with a strong orientational preference. Successful designs were defined as computational predictions with at least a 70% preference for a stable in-plane (deshielding ring interaction) or perpendicular contact (shielding ring interaction) and a distance of less than 6 and 7 Å (5.5 and 6.5 for AF2/AF3 to account for differences between tfmF and Tyr), respectively. Predictions considered “positive” were expected to produce a ring current shift of more than 0.2 ppm (given that most variants without ring currents are dispersed ± 0.2 ppm around the random coil value, **Fig. 1c**), and were subsequently experimentally assessed.

**Figure 2.**
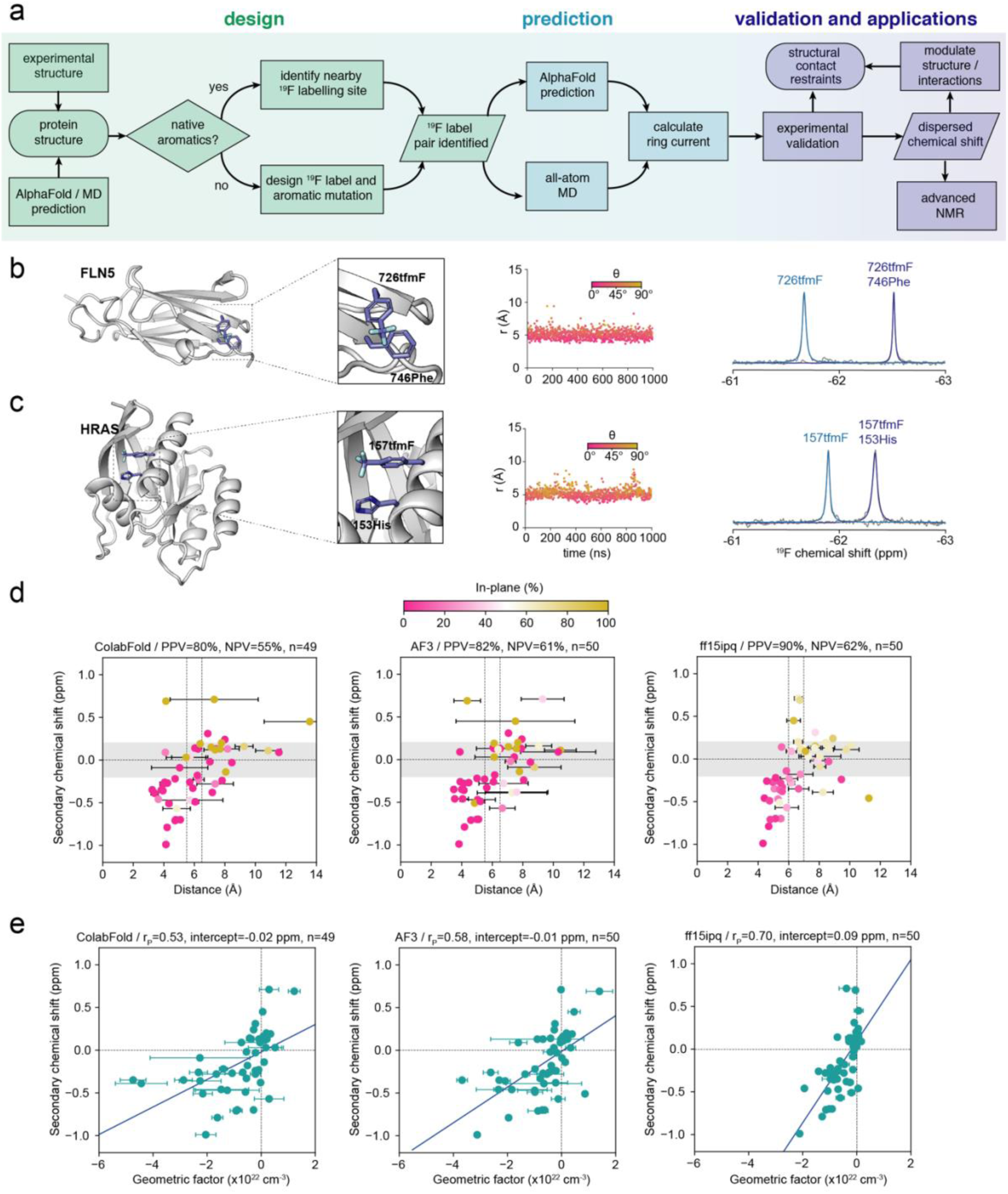
Rational *de novo* design of ring current shifts across different structural motifs. **a.** Flow chart of the design method. **b.** (Left) Structural model of FLN5 726tfmF/746Phe, highlighting an interaction between 726tfmF and 746Phe across two β-strands. (Middle) Distance (*r*) and angle (*θ*) between the CF_3_ group of 726tfmF and aromatic ring of 746Phe observed in a representative all-atom MD simulation. (Right) ^19^F NMR spectra of FLN5 726tfmF with and without 746Phe recorded at 298 K and 500 MHz. **c.** (Left) Structural model of HRAS 157tfmF/153His, highlighting an interaction between 157tfmF and 153His within an α-helix. (Middle) Distance (*r*) and angle (*θ*) between the CF_3_ group of 157tfmF and aromatic ring of 153His observed in a representative all-atom MD simulation. (Right) ^19^F NMR spectra of HRAS 157tfmF with and without 153His recorded at 298 K and 500 MHz. **d.** Scatter plots correlating the distances (coloured by the fraction of time (MD) spent or models (ColabFold/AF3) in the plane of the ring defined as *θ* > 54.6°) with secondary ^19^F chemical shifts for all protein variants predicted by ColabFold (left), AF3 (middle) and MD simulations (right). The error bars represent one s.d. over the five predicted models for ColabFold and AF3, and the s.e.m. obtained from three independent simulations for MD. PPV=positive predictive value; NPV=negative predictive value. Positive secondary chemical shift > 0.2 ppm in magnitude. The vertical lines represent the distance cut-off values for perpendicular and in-plane interactions (lower and higher distance, respectively). **e.** Correlations between predicted geometric factors (1-3cos^2^*θ*)/*r*^3^) and secondary ^19^F chemical shifts, the corresponding Pearson correlation coefficients (r_P_) and intercepts for lines of best fit.

Our design strategy succeeded in predicting proteins with labelling sites engineered to probe a range of structural motifs. For example, the labelling pair within FLN5 726tfmF 746Phe is positioned across two strands of a β-sheet (**Fig. 2b**). Consistent with a shielded ring current predicted by both MD force fields (**Fig. 2b**, **Table S4-S5**) and both ColabFold/AF3 (**Table S3-S4**), its experimental ^19^F chemical shift is −0.70 ppm relative to random coil (**Fig. 2b**). Notably, we found that the same tfmF sidechain in β-sheets can even be designed to contact residues on either adjacent strand (**Fig. S6**). Human HRAS labelled with 157tfmF 153His illustrates the ability to design probes within an α-helix (residues *i* and *i*+4, **Fig. 2c**), while we also engineered successful variants to report on protein tertiary contacts between loops/strands and two different α-helices (HRAS 32tfmF 40Tyr and 137tfmF 94His, **Fig. S5a-f**).

### Geometric descriptors of aromatic interactions predict ^19^F chemical shifts

To evaluate the overall performance of our design method and the extent to which experimental ^19^F chemical shifts can be solely rationalised with ring current effects, we designed and produced a total of 50 different protein variants across six different proteins and one protein complex. These comprised β-sheet proteins FLN5^44^, its neighbouring domain FLN4^45^, titin I27 (27^th^ Ig-like domain of titin)^46^, human filamin A domain 21 (FLNa21)^47^, and the FLNa21 complex with human migfilin^47^, the all α-helical the N-terminal domain (NTD) of *E. coli* HemK^48^, which folds independently of its C-terminal domain^49^, and the mixed α/β human HRAS^50^. For all variants, ColabFold/AF3 predictions and MD simulations with the ff15ipq force field^40^ were performed (see Methods), and the experimental (secondary) ^19^F chemical shifts determined (**Tables S3-S6**). Predictions where ring currents were expected, “negative” variants without ring currents, and also tfmF-labels paired with different aromatic residue types were all experimentally tested to accumulate a diverse dataset.

As designed, the variants generally exhibited a ring current effect at the labelling site whose magnitude correlates with their predicted distances from AlphaFold and MD models (**Fig. 2d**), and with minimal changes to their folding stabilities (**Table S2**). Most of these predictions exhibit a strong preference of tfmF to interact in the perpendicular orientation to the aromatic ring, consistent with nuclear shielding (**Fig. 2d**). Encouragingly, we find that our design strategy has a high success rate (positive predictive value) of 80, 82% and 90% by ColabFold, AF3 and MD simulations respectively (**Fig. 2d, Table S7**). Thus, the false discovery rate is only 10-20%. The prediction of “negative” variants lacking strong ring currents is less robust with a false omission rate of 38-45% (55-62% negative predictive value, **Fig. 2d**, **Table S7**). This is not as crucial for the design strategy, however, as it means some “positive” variants might be missed due to them being predicted as negative variants. When all three prediction methods (ColabFold/AF3/MD) are combined, and a predicted positive design is thus defined as having a ring current by at least one of the prediction tools and a negative when all three methods predict no ring current, then the false omission rate drops to 26% (**Table S7**). Thus, combined use of the prediction tools may be practical to minimise false omissions. Similar statistics were observed with the C36m MD parameters (**Fig. S5g-h, Table S7**). We further note that while labelling/residue pairs that are predicted with ColabFold at moderate to high confidence (pLDDT > 70 and > 90, respectively) for both true and false predictions, AF3 has proportionally fewer moderate-to-high confidence false predictions (**Table S8**), suggesting that AF3-pLDDT values may be useful in prioritising design candidates.

We also evaluated the correlation between the secondary ^19^F chemical shifts and a geometric term quantifying the ring current effect, (1-3cos^2^*θ*)/*r*^3^, where *r* and *θ* are the distance and angle of the CF_3_ group in tfmF relative to the aromatic ring^51^, respectively (**Fig. 1e, 2e**). These analyses show moderately positive correlations (Pearson correlation coefficient, r_P_) of 0.53 and 0.58 for ColabFold and AF3, respectively. Thus, although AlphaFold predictions perform well in predicting good variants they are limited in “ranking” variants according to the magnitude of the ring current-induced secondary chemical shift. MD simulations, however, showed a stronger r_P_ of 0.7 highlighting that these predictions are more reliable in ranking designed variants (**Fig. 2e**). More importantly, these results show that ring current descriptors predict secondary ^19^F chemical shifts remarkably well (i.e., ∼50% of the variation in the experimental data is explained by ring currents). The proteins simulated with both force fields (see Methods) show MD simulations exhibiting correlations of up to 0.8 (64% of variation explained, **Fig. S5h**). The unaccounted variation by our ring current model therefore has likely contributions from other factors such as Van der Waals interactions and electrostatics^24^, in line with our observations that removing aromatic rings by mutagenesis reduces the magnitude of the secondary chemical shift by ∼50-80% (**Figs. 1c, 2b-c, S6c**). Overall, our approach enables the rational design of proteins with ^19^F-labelling sites, whose chemical shifts are engineered to probe distance- and angular-dependent sidechain contacts.

### Discovery of alternative conformational states using ring current design

We next explored whether our ring current design strategy could be employed to elucidate alternative conformational states of proteins, where predictive strategies such as AlphaFold often fail. We examined the *E. coli* N^5^-glutamine methyltransferase HemK N-terminal domain (NTD), which is composed of a five-helix bundle^48^. We designed HemK NTD tfmF-labelled at position 38 to report on the structure of helix 3 by contacting Phe42 (an (*i*, *i*+4) contact, **Fig. 3a-b**). The ^19^F NMR spectrum of HemK 38tfmF indeed showed shielding relative to random coil, and the Phe42Ala substitution confirmed the chemical shift is induced by a ring current effect (**Fig. 3c**). As a proof-of-concept, we examined an existing variant in the lab with mutations within and around helix 3 (mutant, I26V/R34K/Q46R). While ColabFold and AF3 models predict no structural changes (**Fig. S7**), 38tfmF-labelling of the mutant resulted in a random coil ^19^F chemical shift, indicating the loss of the local ring current (**Fig. 3c**), despite ^1^H,^15^N-correlated NMR spectra showing that both variants were globally folded (**Fig. S8**). We performed additional long-timescale MD simulations of wild-type and mutant HemK (see Methods). Six independent simulations, each lasting 20 μs, showed that the protein backbone exhibits increased flexibility in the h3 region of mutant HemK relative to that of wild-type (**Fig. 3d**), due to local helical unfolding (**Fig. 3e, S9)**. The ^19^F design therefore provides a simple and fast approach to afford structural insights into mutation-induced conformational changes without the need for multi-dimensional spectral assignment.

**Figure 3.**
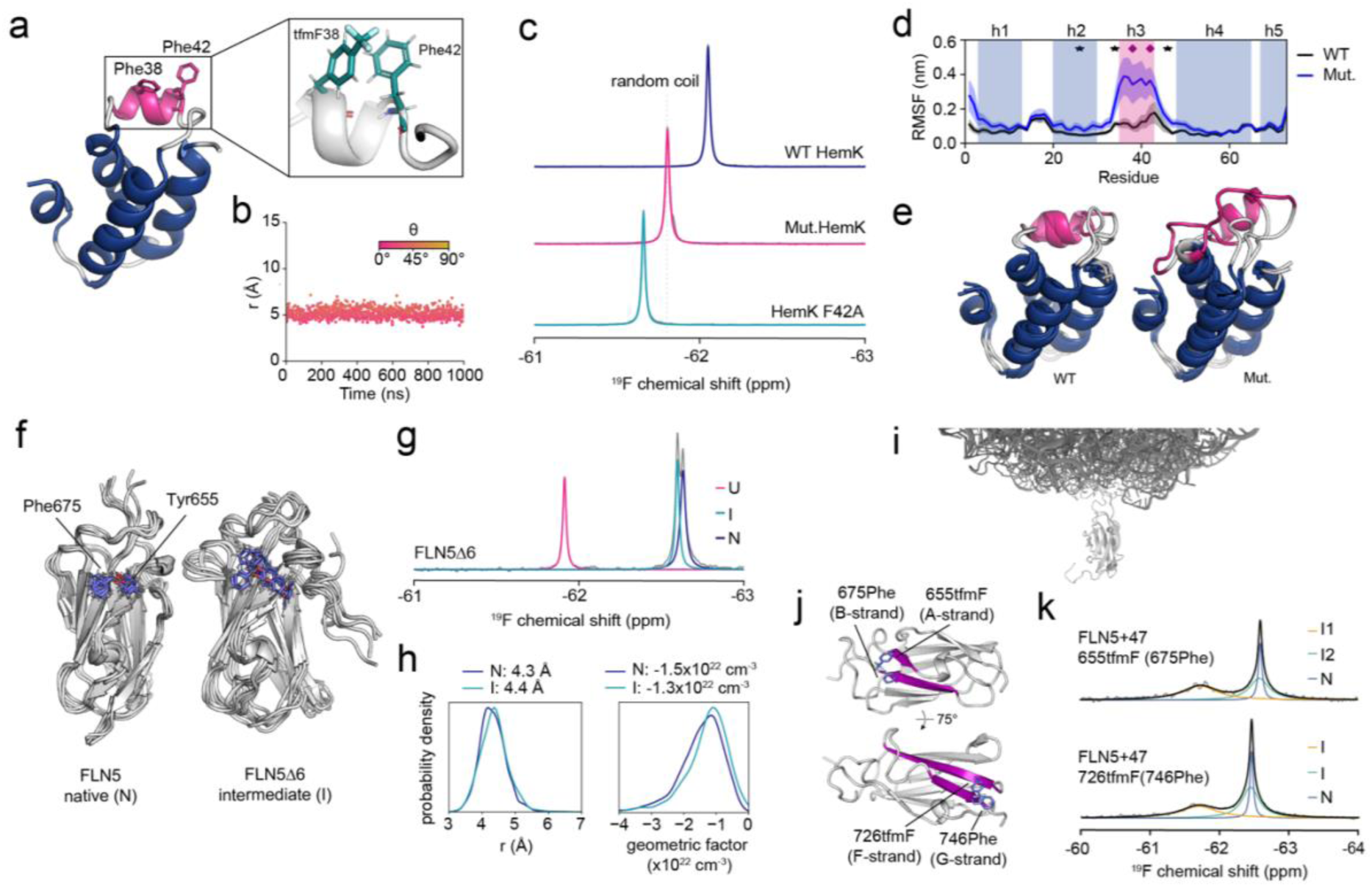
Engineered ring current shifts enable detection and characterisation of alternative protein conformational states. **a.** Crystal structure (left) of the HemK NTD (residues 1-73, PDB 1T43) and predicted interaction between 38tfmF and Phe42 observed in all-atom MD simulations (right). **b.** Distance (*r*) and angle (*θ*) between the CF_3_ group of 38tfmF and aromatic ring of Phe42 observed in a representative all-atom MD simulation. **c.** ^19^F NMR spectra of HemK NTD 38tfmF variants (mutant containing I26V/R34K/Q46R mutations) and the Phe42Ala mutant, recorded at 298 K and 500 MHz. **d.** Backbone (Cα) root mean square fluctuation (RMSF) analysis for HemK NTD observed in long-timescale all-atom MD simulations (average ± s.e.m. from six independent simulations of 20 μs). The stars indicate the mutation locations, and the diamonds represent the tfmF labelling site and aromatic ring location within helix h3. **e.** Representative structures of wild-type and mutant HemK obtained from all-atom MD simulations. **f.** Structural ensembles of natively folded (N) FLN5 (left) obtained from an all-atom MD simulation with the ff15ipq force field and FLN5Δ6 folding intermediate (I)^33^. The Tyr655 and Phe675 sidechains are shown as sticks for both ensembles. **g.** ^19^F NMR spectrum of FLN5Δ6 655tfmF recorded at 283 K and 500 MHz^8,9^ showing unfolded (U), intermediate (I) and native (N) conformations at equilibrium. **h.** Probability distributions of *r* and the geometric factor (1-3cos^2^*θ*)/*r*^3^) calculated for the FLN5 N and I state and average values. **i)** Structural model of a folded FLN5 nascent chain tethered to the ribosome **j)** Structural models of FLN5 655tfmF/675Phe and 726tfmF/746Phe highlighting the A/B and F/G strand pairs, respectively. **k)** ^19^F NMR spectra of FLN5+47 RNCs with the 655tfmF/675Phe and 726tfmF/746Phe labelling pairs, recorded at 298 K and 500 MHz. For FLN5+47 655tfmF/675Phe two folding intermediates (I1 and I2) have been identified previously^8^.

We next explored applications to protein folding intermediates. We have previously determined the structural ensemble of a high-energy folding intermediate of FLN5Δ6^33^, where the six C-terminal (G-strand) residues of full-length FLN5 are truncated. This intermediate (I) exhibits increased flexibility (relative to the native state, N) with a disordered C-terminus, while the remaining strands have a native-like conformation (**Fig. 3f**). We used the 655tfmF labelling site which experiences a ring current from Phe675 (**Fig. 1c-f**) to test whether the contacts between the A- and B-strands indeed remain native. The ^19^F NMR agrees with the structural model and shows that the I state has a near-native chemical shift, strongly shielded relative to the unfolded (U) state (**Fig. 3g**). The small reduction in shielding observed for the I state may be accounted for by increased protein dynamics resulting in slightly reduced interactions with the aromatic ring (**Fig. 3h**).

Having established that ring currents measured by ^19^F NMR can be used to characterise partially folded protein conformations with two simple model proteins, we set out to tackle a more challenging system. Protein folding begins during biosynthesis on the ∼2.5-MDa ribosome (**Fig. 3i**), where co-translational folding (coTF) intermediates have been found to be thermodynamically stabilised (relative to off the ribosome) by up to 5 kcal mol^-1^ (ref.^8,9^). Such ribosome-bound intermediates have not been directly observable by cryo-electron microscopy due to their dynamic nature, nor by NMR using ^15^N or perdeuteration combined with selective ^1^H,^13^C-methyl labelling due to fast transverse relaxation rates induced by even transient ribosome interactions^9,34^. Direct observations have, however, been made using more sensitive ^19^F NMR measurements^8^, although details of the folding pathway(s) have remained elusive^8,9^. We have previously shown that FLN5 folds co-translationally via two folding intermediates by tfmF-labelling residue 655, revealing one intermediate with a random coil chemical shift (I1) and one intermediate with a native-like chemical shift (I2)^8^. These chemical shifts can now be attributed to the absence and presence of the Tyr655-Phe675 contact (**Fig. 1c**) and thus likely an unfolded A-strand and native A-B strand contacts, respectively (**Fig. 3j-k**). Previous attempts to further resolve I1 and I2 on the ribosome with other labelling sites have, however, failed due to substantial line broadening and poor dispersion distinguishing native and non-native (random coil) chemical shifts^8^.

We previously hypothesised that the C-terminal G-strand may be unfolded in the coTF intermediates^8^, similar to the folding intermediate of FLN5Δ6 (**Fig. 3f**). We therefore used our 726tfmF/746Phe engineered ring current (**Fig. 2b**), whose large secondary chemical shift directly reports on interactions between the F- and G-strands. By labelling ribosome-bound nascent chain complexes with this labelling pair, we identified two broad resonances attributable to intermediate states and possessing a random coil and native-like 726tfmF/746Phe chemical shift (**Fig. 3j-k**), confirming that one coTF intermediate has indeed an unfolded G-strand. This example illustrates the power of our design approach to study proteins part of MDa-sized biomolecular complexes that elude other structural biology methods.

### Detection and structural characterisation of protein-ligand interactions

^19^F NMR is a common tool used in drug discovery and characterising protein-ligand interactions^14^. We tested whether ligand binding to proteins could be detected and structurally interpreted with our method. Noticing that residue 28 of human HRAS points into the ligand (GDP) binding site (**Fig. 4a-b**), we modelled the structure and dynamics of the HRAS-GDP complex 28tfmF using MD and found that 28tfmF is predicted to be in-range to experience a ring current from the guanine ring system (**Fig. 4c**, **Table S5**). To verify this experimentally, we purified recombinant HRAS expressed in *E. coli* and found that in the presence of GDP/Mg^2+^ HRAS 28tfmF exhibits a single peak shielded (−62.24 ppm) relative to random coil (**Fig. 4d**). In the absence of GDP/Mg^2+^ there was an additional peak at −61.78 ppm corresponding to the apo protein. The residual amount of GDP-bound HRAS is likely due to the picomolar affinity of GDP for HRAS^52^. Ring currents can thus be used to both directly detect and structurally characterise and validate protein-ligand binding poses.

**Figure 4.**
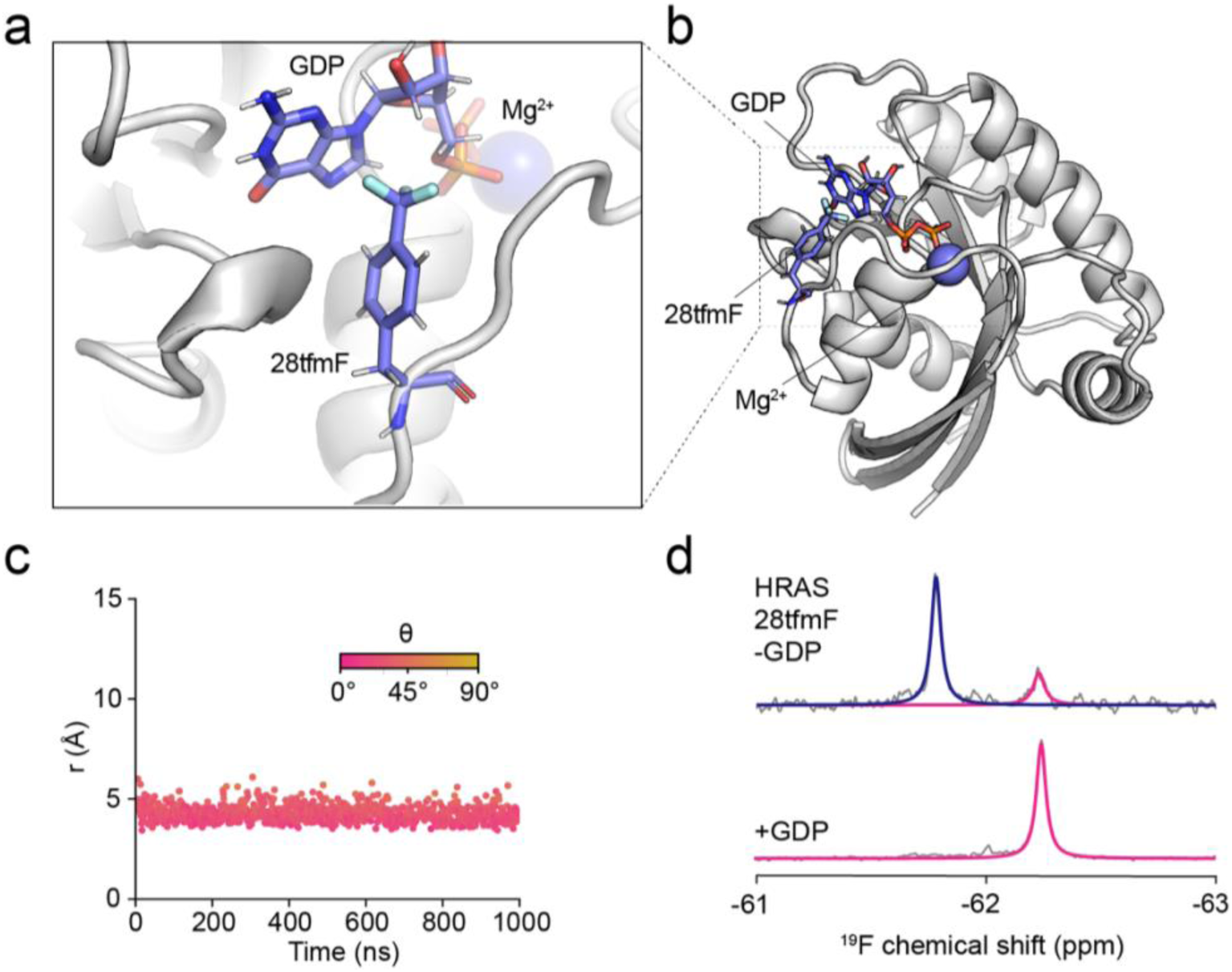
Probing protein-ligand interactions. **a-b.** Structural model of HRAS 28tfmF bound to GDP obtained from all-atom MD simulations. **c.** Distance (*r*) and angle (*θ*) between the CF_3_ group of 28tfmF and aromatic rings of GDP observed in a representative all-atom MD simulation. **d.** ^19^F NMR spectrum of HRAS 28tfmF purified in the absence of Mg^2+^/GDP (top) and presence of Mg^2+^/GDP (bottom) recorded at 298 K and 500 MHz.

### Protein folding thermodynamics determined in cells

We next turned to explore novel applications of our approach in cells. The crowded cellular environment has been shown to influence the interactions, structures, and folding thermodynamics of proteins^11,53,54^. Well-resolved intracellular NMR resonances are, however, limiting the application of in-cell NMR to a handful of small protein domains. We first sought to examine how the folding thermodynamics of FLN5 is altered in vivo, and thus used the destabilising Phe672Ala mutant to enable population of both the unfolded and folded states at equilibrium^9^. However, natively folded FLN5 resonances are undetectable in-cell by ^1^H,^15^N-correlated NMR^55^, and conversely, intracellular unfolded cross-peaks are severely overlapped using selective ^1^H/^13^C-methyl labelling (**Fig. S10**). Yet despite the broad linewidths of the in-cell spectra (up to ∼400 Hz vs 10 Hz of purified protein), using ^19^F NMR and owing to the large secondary chemical shift induced by the ring current between the 655tfmF-labelling site and Phe675 ring, we are able to detect, resolve, and quantify the two conformational populations (**Fig. 5a**) and their equilibrium stabilities to extract the thermodynamic parameters of folding in cells (**Figs. 5d, S11**), and compare them to a cell lysate sample (**Fig. 5b**) and purified protein (**Fig. 5c**).

**Figure 5.**
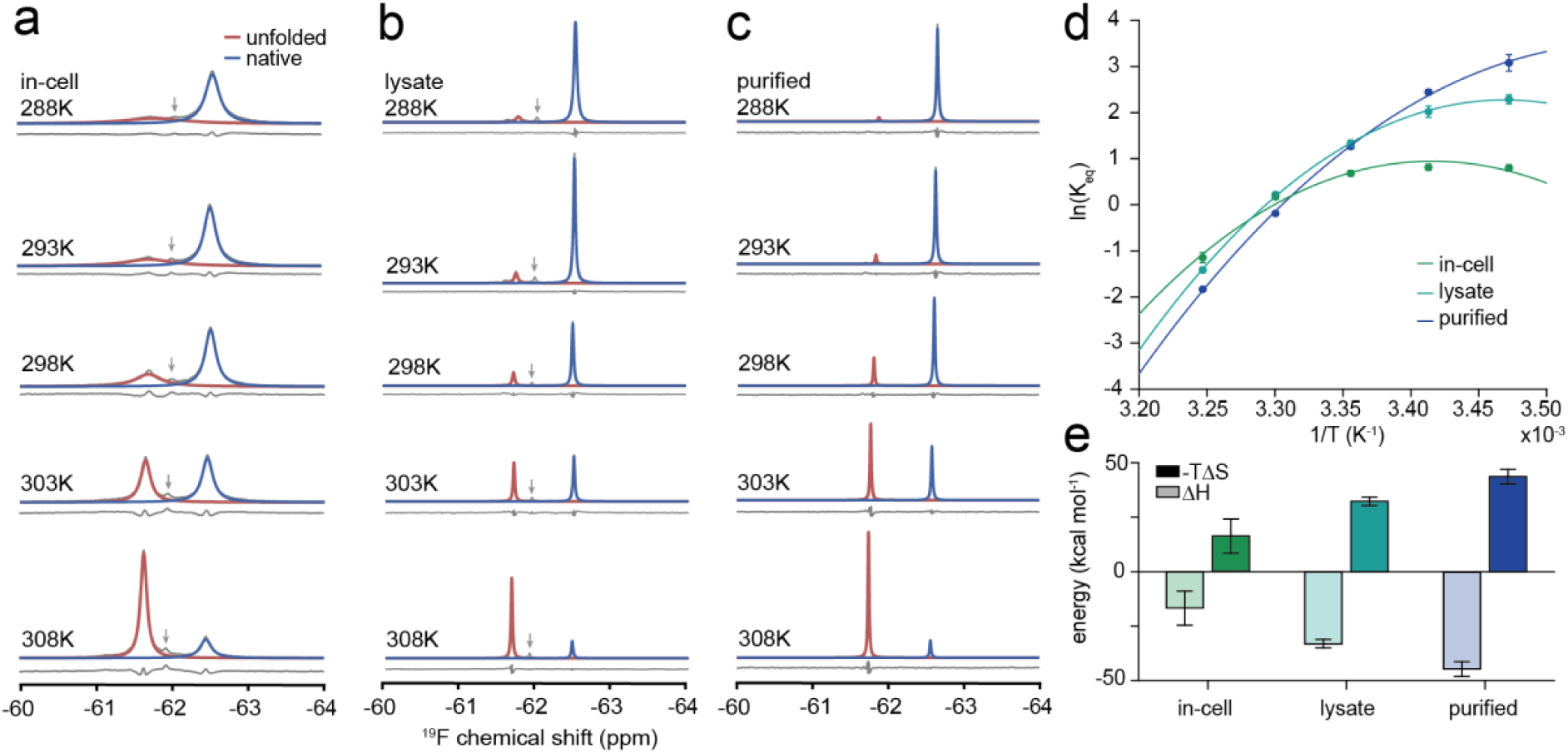
Quantitative protein thermodynamics measured inside living cells. **a.** In-cell, **b.** in-lysate and **c.** purified ^19^F NMR spectra of FLN5 672Ala^9^ at various temperatures recorded at 500 MHz showing an unfolded and folded population. The arrow indicates the position of the free tfmF peak. **d.** Temperature-dependence of the folding equilibrium constant (K_eq_) in all conditions. Data were fit to a modified Gibbs-Helmholtz equation^9^. Error bars represent the propagated s.e.m. from NMR lineshape fits obtained by bootstrapping. **e.** Thermodynamic parameters obtained from the fits in panel d (mean ± 95% CI of fitted parameters).

The temperature-dependence of folding is reduced in cells relative to buffer and the stability of the protein altered. At low temperatures, we find that FLN5 is destabilised relative to the purified sample (ΔΔG_folding,cell-buffer_ = +1.4 ± 0.1 kcal mol^-1^ at 288 K) whereas at higher temperatures (and physiological temperature) the protein is stabilised in cells (ΔΔG_folding,cell-buffer_ = −0.4 ± 0.1 kcal mol^-1^ at 308 K). The fitted thermodynamic parameters reveal that the enthalpy of folding becomes less negative in cells, and the entropic penalty of folding is reduced (ΔΔH _folding,cell-buffer_ = +27.8 ± 8.5 kcal mol^-1^;-TΔΔS _folding,cell-buffer_ = −27.5 ± 8.5 kcal mol^-1^ at 298 K), an observation that was also found for nascent proteins folding during translation on the ribosome^9^. This is likely due to electrostatic interactions between the unfolded state and other molecules in the cell^11,56^ dominating at lower temperature and entropic excluded volume effects exerted on unfolded polypeptides^56^ at higher temperatures. Thus, in cells, quinary interactions of proteins affect their thermodynamic landscape and energetic factors underlying fundamental biophysical processes such as folding, in line with previous reports of other proteins^11,53,54,57^. These data suggest that further reductions in the entropic penalty of co-translational protein folding^9^ occur in the cellular environment, and folding could thus be entropically driven on the ribosome *in vivo*.

### Structural validation of protein-protein interaction modes in cells

Lastly, we studied the human FLNa21-migfilin complex to explore in-cell protein-protein interactions. Migfilin is a disordered adaptor protein that binds to the FLNa21 filamin, forming an additional strand to one of its β-sheets^47^ (**Fig. 6a**). The published crystal structure of the complex contains two FLNa21 molecules with migfilin sandwiched between the folded domains, and *in vitro* NMR data suggests that the interaction with FLNa21 chain A is the dominant interaction in solution^47^. We therefore focused on designing a ring current labelling pair for this interaction mode to validate the structural model in cells. The crystal structure reveals that Phe14 in migfilin could induce a ring current to potential FLNa21 labelling sites on the C-strand (**Fig. 6a**). MD simulations predicted, however, that regardless of the labelling site (2274tfmF and 2272tfmF) on FLNa21 there is no strong ring current interaction between these sidechains (**Figs. 6b, S12, Tables S5-S6**); indeed, titrations of migfilin into FLNa21 2274tfmF *in vitro* showed only a small ^19^F chemical shift difference between the bound and free form of ∼0.03 ppm, and which could not be resolved by lineshape analysis. MD simulations predicted, however, that the migfilin Ser12His mutant produces a strong ring current interaction with 2274tfmF (**Figs. 6c-d, S12, Tables S5-S6**). This was confirmed experimentally with a titration *in vitro* showing the ^19^F chemical shift of the bound state is shielded by −0.49 ppm (**Fig. 6e**), and where the increased chemical shift dispersion permits lineshape analysis and accurate quantification of the binding affinity, found to be in the low micromolar range (**Fig. S13c**) in line with previous work^47^.

**Figure 6.**
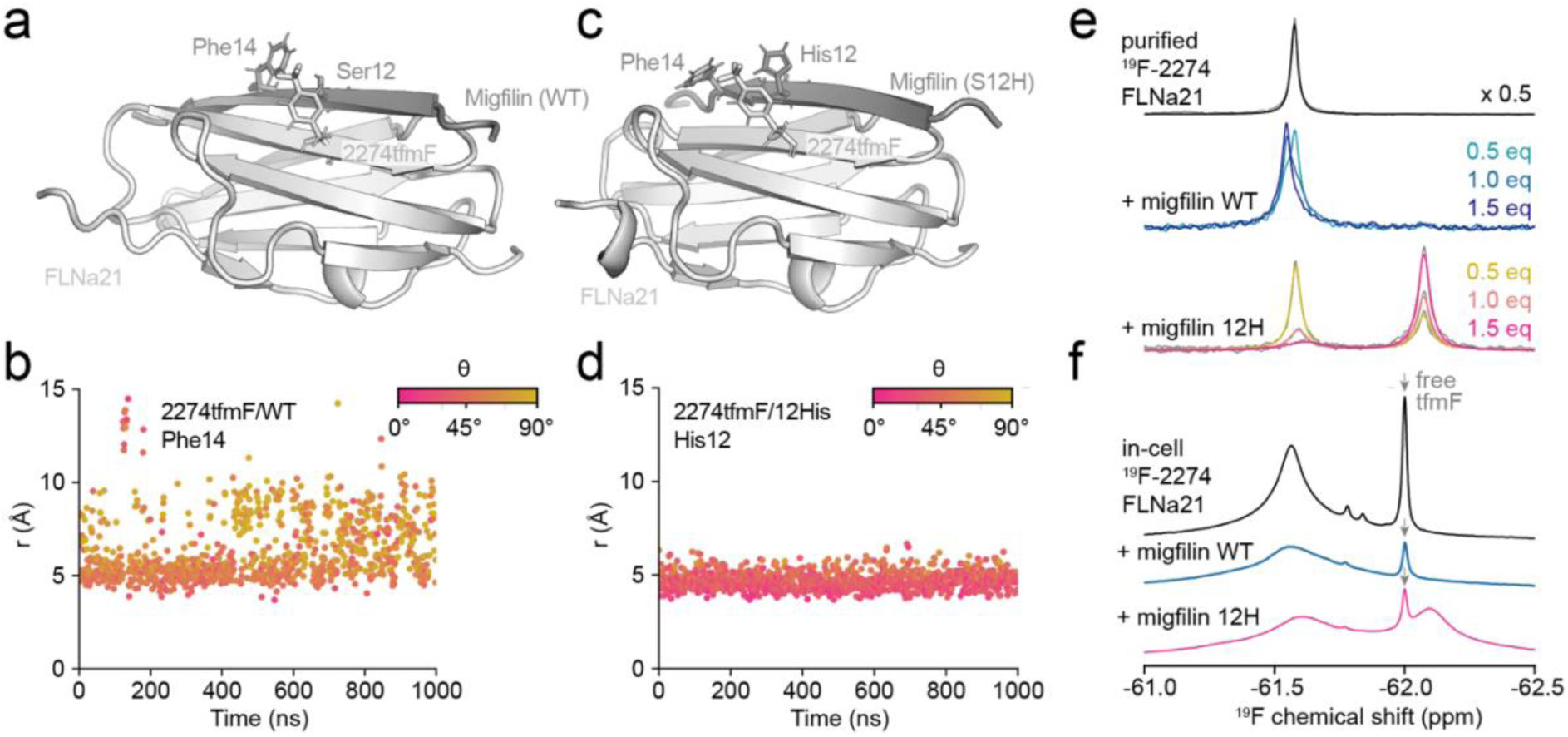
Direct and residue-specific detection of protein-protein interactions in cells. **a.** Structural model of the FLNa21 (2274tfmF) – migfilin (WT) complex obtained from all-atom MD simulations. **b.** Distance (*r*) and angle (*θ*) between the CF_3_ group of 2274tfmF and the aromatic ring of Phe14 observed in a representative all-atom MD simulation. **c.** Structural model of the FLNa21 (2274tfmF) – migfilin (Ser12His) complex obtained from all-atom MD simulations. **d.** Distance (*r*) and angle (*θ*) between the CF_3_ group of 2274tfmF and the aromatic ring of 12His observed in a representative all-atom MD simulation. **e.** ^19^F NMR spectrum of purified FLNa21 2274tfmF alone and with 0.5-1.5 molar equivalents (eq) of migfilin WT and 12His. **f.** In-cell ^19^F NMR spectrum of FLNa21 2274tfmF and co-expressed with WT and 12H migfilin. All spectra were recorded at 298 K and 500 MHz.

We then explored in-cell NMR of FLNa21-migfilin binding, where FLNa21 was expressed as a GST-fusion protein^47^. 2D ^1^H,^15^N-SOFAST HMQC spectra of in-cell FLNa21 showed no detectable resonances of the domain but resonances could be recovered upon lysis (**Fig. S13a**), showing that traditional in-cell NMR of this system is intractable. While ^19^F in-cell NMR of FLNa21 2274tfmF enabled detection of GST-FLNa21 in *E. coli*, when co-expressed with WT-migfilin, the bound complex could not be discerned unambiguously due to substantial line broadening paired with a limited chemical shift difference with the free state (**Fig. 6f**). When migfilin 12His is instead co-expressed, however, we clearly observe an additional ^19^F NMR peak coinciding with the bound form as observed *in vitro* (**Figs. 6f, S13b**). This resonance unambiguously confirms the protein-protein interaction in the crowded intracellular environment and validates the *in vitro* structural model previously determined^47^.

## Discussion

We have presented a rational protein design strategy to yield structurally interpretable ^19^F chemical shifts with improved chemical shift dispersions. The approach is relatively straightforward, requiring only the introduction of a commercially available ^19^F-label near native or engineered aromatic residues, without the implementation of new pulse-sequences nor chemical synthesis. The design can be guided most conveniently and rapidly by AlphaFold at a high success rate (>80%), with reduced false omissions and higher accuracies achieved by MD simulations, where non-canonical ^19^F-labelled amino acids can be explicitly modelled. Our method was validated both experimentally, by comparison of tfmF-labelled proteins with and without the accompanying aromatic residue, and by correlating the predicted magnitude of the ring current effect with the observed secondary chemical shifts (**Fig. 2**). The applications in this work demonstrate its utility in determining how modulations (mutations, ribosome-bound conformations, ligand-unbinding, in-cell environment) affect protein structure and energetics, and future applications may include validation of novel, predicted structural states, for example, from MD simulations.

Our dataset illustrates that aromatic ring current effects dominate modulations to the chemical shift of solvent-exposed ^19^F NMR labelling sites of cytosolic proteins (**Fig. 2d-e**), and which is exploited in our labelling method. The chemical shift of the engineered labels can therefore be interpreted as a residue pair-specific Van der Waals contact with an upper bound distance between two amino acids, generally <7 Å for any ring-current, or a more precise distance determined by MD, where longer simulations and/or enhanced sampling methods^58^ provide improved precision. The information extracted from ring current contacts complements and resembles interpretation of other types of experimental data probing short-range contacts, such as nuclear Overhauser effects (NOEs), recently developed optical measurements of intramolecular protein distances^59^, and longer-range measurements from (^19^F) paramagnetic relaxation enhancement (PRE) NMR, cross-linking mass spectrometry (XL-MS), FRET, and DEER. Upper bound distances of residue pairs, explicit modelling or *post hoc* rotamer calculations may be used to restrain structural models, similar to approaches employed to calculate FRET^60^ and PRE/DEER^61^ data. Ring current contact data from ^19^F NMR could thus be applied as restraints in modelling using MD simulations^62^ and directly in AlphaFold-based pipelines as recently illustrated with XL data^63,64^. Further exploitation of our dataset may also enable more detailed chemical shift prediction methods to account for other effects such as electrostatics^24,65^. Its applicability to predicting ^19^F chemical shifts of buried amino acids may be limited, however.

We have tested our labelling strategy across 6 proteins with different folds. However, a potential limitation of our dataset in assessing correlations between chemical shifts and structure is that β-strand proteins and ^19^F-ring contacts across two β-strands are overrepresented relative to contacts within helices and between different secondary structure elements. Our dataset contains predominantly shielding ring current effects (interactions perpendicular to the ring plane). This may to some extent also reflect the intrinsic propensity of tfmF to interact with aromatic rings via π-π stacking interactions for labelling pairs within β-sheets (**Fig. 2b**) and α-helices (**Fig. 2c**). The performance of our method, and indeed the (de)stabilising nature of ^19^F-labelling, may therefore vary depending on the target protein, structural element and unnatural amino acid used.

While tfmF is advantageous due to its low number of rotatable sidechain dihedral angles compared to BTFMA (**Fig. S3**), we certainly expect that our strategy may be extended to other fluorinated amino acids in the future, such as recently developed pyridone-based^16^ and monofluorinated^3^ probes. Our approach is also independent of their incorporation methods, whether genetically encoded, by selective pressure incorporation, or by covalent conjugation. In this work, we have focused on the use of phenylalanine and histidine as engineered ring current sources due their smaller sizes compared to tyrosine and tryptophan, although exemplar variants show these residues do not preclude induction of ring currents (e.g., native Trp2262 in FLNa21 tfmF2242 and Tyr40 in HRAS 32tfmF, **Table S5**).

We have showcased the capabilities of our method by examining a range of different biological systems. Our exemplar applications included the identification of alternative conformational states, the characterisation of nascent protein folding intermediate structures on the ribosome, and probing protein-ligand and protein-protein interactions; each of which have permitted structural characterisation or validation, and where ^19^F NMR is the only available experimental method capable of directly obtaining residue-resolution data of co-translational folding intermediates outside the ribosome exit tunnel^8,9^. Our method also overcomes challenges with spectral overlap associated with CF_3_-probes, and this has permitted in-cell NMR measurements to validate *in vitro* structural models and quantify thermodynamics in the cell.

Rationally designed fluorinated proteins together with simple 1D pulse-acquire experiments permit access to structural characterisations of complex biological systems, where multi-dimensional ^19^F NMR methods can suffer from sensitivity limitations^1,27^. We anticipate future applications of more advanced NMR experiments will include, amongst other measurements, those of protein dynamics across different time-scales; for example, CEST experiments^8^ will particularly benefit from the increased spectral dispersion of chemical shifts of different conformational states. The rational design of ^19^F-labelling sites will therefore support new technical capacities and biological applications of NMR spectroscopy.

## Methods

### Molecular biology and protein production

All mutations (including amber stop codons) were introduced using standard site-directed mutagenesis procedures. FLN5, FLN4, titin I27, HemK NTD (residues 1-73), and HRAS were expressed and purified as hexahistidine-tagged constructs as previously described^8,9,34^. Expression of GST-tagged FLNa21^47^ was performed as for the other proteins and purified using a similar procedure but replacing the Ni^2+^-NTA resin with a Glutathione Sepharose^TM^ 4B resin (Cytiva^TM^). Elution was achieved in the presence of 10 mM reduced L-glutathione. Migfilin (residues 1-85) DNA constructs and production were previously described^47^. Labelling with 4-trifluoromethyl-L-phenylalanine (tfmF) was achieved by amber suppression: cells were co-transformed with the pEVOL-pCNF-RS suppressor plasmid^66,67^, and expression of the orthogonal pair induced by addition of L-arabinose (0.2% (w/v)), as previously described^8^.

### NMR spectroscopy

NMR data were acquired on 500- and 800-MHz Bruker Avance III spectrometers, both equipped with TCl cryoprobes, and all recorded using TopSpin3.5pl2 at 298 K unless stated otherwise. All samples were measured in Tico buffer (10 mM HEPES, 30mM NH_4_Cl, 12mM MgCl_2_, 2mM β-merceptoethanol, pH 7.5) supplemented with 10% (v/v) D_2_O and 0.001% (w/v) sodium trimethylsilylpropanesulfonate (DSS) as the reference compound. 1D ^19^F pulse-acquire experiments were recorded with an acquisition time of 350 ms and a recycle delay of 1.5 s. To monitor the stability of the FLN5+47 726tfmF/746Phe RNC, short successive spectra were recorded, such that data with detectable changes over time (from nascent chain release or degradation) were discarded, and only data from intact RNC complexes were summed together, as previously described^9,34^. 2D ^1^H, ^15^N SOFAST-HMQC experiments were performed with direct and indirect dimension acquisition times of 50.4 and 295 ms, respectively, and a recycle delay of 100 ms. Data processing and analysis were performed with nmrPipe^68^, CCPN Analysis^69^, MATLAB (R2014b, The MathWorks Inc.) and Julia^70^ as previously described^8^. The time-domain ^19^F NMR data were multiplied with an exponential window function with a line broadening factor of 5-10 Hz prior to Fourier transformation, and subsequently baseline corrected, and line shapes analysed using Lorentzian functions^8^.

### In-cell NMR spectroscopy

High-density *E. coli* cell cultures for in-cell NMR were prepared using the protocol for production of purified RNC samples as previously described^8,9,34,71^. Protein expression was induced with 1 mM IPTG (0.25 mM for FLNa21+migfilin cell cultures where both proteins were induced with IPTG) for ∼16 h at 30°C. Cells were then harvested by centrifugation at 4,000 rpm for 15 min, washed three times with and subsequently resuspended in in-cell NMR buffer (75 mM bis-tris propane, 75 mM HEPES, 25 mM MgCl_2_, pH 7.5). Cells slurries were prepared at a concentration of 40% (w/v) and supplemented with 10% D_2_O and 0.001% (v/v) DSS. To monitor sample stability, we recorded short spectra in succession to identify spectral changes over time; no discernible changes were observed between spectra acquired during the course of NMR data acquisition. Cell leakage of ^15^N-labeled samples was monitored using 1D ^1^H,^15^N-correlated SORDID diffusion measurements interleaved between 2D SOFAST-HMQC experiments and acquired with a diffusion delay of 300 ms, using gradient pulses of 4 ms and gradient strengths of 5% and 95% of the maximum gradient strength (0.56 Tm^-1^)^55,72^. Given the sensitivity limitations of ^19^F diffusion measurements, we instead assessed cell leakage of ^19^F-labelled samples by recording equivalent NMR experiments of the supernatant after centrifugation at 9000 rpm for 10 min. Acquisition parameters for all experiments were identical to those performed on purified samples described above. For the quantification of protein folding thermodynamics, we used MATLAB to fit the temperature-dependent folding equilibrium constant, *K_eq_* (directly measured from peak integrals), to a modified Gibbs-Helmholtz equation to determine the change in entropy (Δ*S*), enthalpy (Δ*H*) and heat capacity of folding (Δ*C*_p_) of folding:

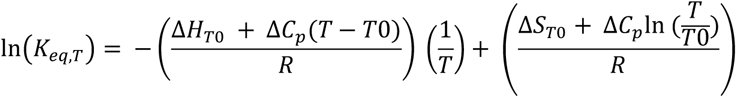

*T* and *T_0_* are the temperature of the measurements and standard temperature (298 K), respectively. We report the errors of the fitted parameters as the 95% confidence interval (CI).

### MD simulations of fluorinated protein variants

All MD simulations were performed using GROMACS version 2021 or 2023^73^. Independent simulation replicates were initiated from different sidechain rotamers of either the fluorinated amino acid and/or nearby aromatic residue where possible. Proteins listed in Table 1 (and their fluorinated variants) were parametrised using the CHARMM36m (C36m) force field^41^, including parameters for 4-trifluoromethyl-L-phenylalanine (tfmF, July 2021 release of the C36m force field files for GROMACS)^42^. Simulations were initiated from the following experimental structures, after introducing tfmF and the relevant mutations *in silico* using PyMol (version 2.3, Schrödinger, LLC). The C36m force field resulted in partial unfolding of wild-type HemK and HRAS, which were therefore not further simulated with C36m. We thus describe the analysis of all proteins (with ff15ipq, see below) in the main text and a common dataset comparison with C36m in the Supplementary Information.

**Table 1.** List of proteins simulated with the C36m force field including their PDB/AlphaFold2 database codes and references. The FLN5 (650-660) peptide was used as a reference to calculate the solvent accessibility of the CF_3_ group in tfmF for an unfolded peptide.

| Protein (residues) | Structure code | Source | Reference |
| --- | --- | --- | --- |
| FLN5 (650-660)<br>peptide | N/A | Linear peptide built in<br>PyMol | N/A |
| FLN5 (645-750) | 1QFH/6G4A | X-ray<br>crystallography/NMR | 33,44 |
| FLN4 (547-648) | AF-P13466-F1 | AlphaFold2 | 28,74 |
| FLNa21 (2236-2328) | 2W0P | X-ray crystallography | 47 |
| FLNa21 (2236-2328) /<br>Migfilin (8-16) complex | 2W0P | X-ray crystallography | 47 |
| I27 (1-89) | 1TIT | NMR | 46 |

Protein molecules were placed in the centre of a dodecahedral-shaped box at least 1.2 nm away from the box edge. The systems were then solvated and MgCl_2_ was added at a final concentration of 12 mM to neutralise the simulation box. Energy minimisation was run using the steepest descent algorithm. Dynamics simulations then employed the leap-frog integrator and a 2 fs timestep along with the LINCS algorithm^75^ to constrain all bonds connected to hydrogen. Nonbonded interactions were cut off at 1.2 nm, and an additional switching function between 1.0 and 1.2 nm was applied to Lennard-Jones interactions. The Particle Mesh Ewald (PME)^76^ method was used for long-range Coulombic interactions with cubic interpolation and a fourierspacing of 0.16 nm. The temperature was kept constant using the velocity rescaling algorithm^77^ with a time constant of 0.1 ps. The simulation systems were then first equilibrated in the NVT ensemble for 500 ps at 298 K using position restraints on all heavy atoms (1000 kJ mol^-1^ nm^2^ along all coordinate axes), followed by an additional 500 ps in the NPT ensemble at the same temperature and with the same restraints. Pressure was controlled using the Berendsen barostat^78^, set to 1 bar with a compressibility of 4.5×10^-5^ bar^-1^ and coupling constant of 2 ps. Production simulations in the absence of any restraints were then run for 1 μs in the NPT ensemble using the Parrinello-Rahman algorithm^79^, saving protein coordinates every 0.1 ns.

The following proteins in Table 2 were simulated using the AMBER ff15ipq force field^39^ and SPC/E_b_ water parameters^80^. Parameters for tfmF were taken from ref.^40^. Systems were prepared using *tleap* from AmberTools23^81^. Proteins were placed in a truncated octahedral box and neutralised with sodium ions. The ParmEd python library^82^ was then used to convert the AMBER files to GROMACS format. Simulations were conducted using a 1.0 nm cut-off distance for nonbonded interactions and the PME method^76^ for long-range Coulombic interactions with cubic interpolation and a fourierspacing of 0.125 nm. The remaining run parameters and protocols were identical to the C36m simulations described above. As mentioned above, HemK and HRAS (in complex with Mg^2+^-GDP) were only simulated with ff15ipq parameters as we observed protein instability during microsecond timescale MD simulations with C36m for these systems.

**Table 2.** List of proteins simulated with the ff15ipq force field including their PDB/AlphaFold2 database codes and references.

| Protein (residues) | Structure code | Source | Reference |
| --- | --- | --- | --- |
| FLN5 (645-750) | 1QFH/6G4A | X-ray crystallography/NMR | 33,44 |
| HemK (1-73) | 1T43 | X-ray crystallography | 48 |
| FLN4 (547-648) | AF-P13466-F1 | AlphaFold2 | 28,74 |
| FLNa21 (2236-2328) | 2W0P | X-ray crystallography | 47 |
| FLNa21 (2236-2328) / Migfilin (8-16) complex | 2W0P | X-ray crystallography | 47 |
| I27 (1-89) | 1TIT | NMR | 46 |
| HRAS (1-166, Mg<sup>2+</sup>-GDP) | 4Q21 | X-ray crystallography | 50 |

### GDP parameterisation

For the HRAS complex, GDP bonded and Lennard-Jones parameters were assigned based on the GAFF2 force field^83^ with *antechamber*^84^. Using AM1-BCC partial charges^85^ as a starting point, we then parameterised implicitly polarised charges (IPolQ)^86^ to obtain GDP charges that are compatible with the ff15ipq force field^39^. A protocol described in the following github repository was followed: https://github.com/darianyang/ff15ipq-lig. AMBER20^87^ and ORCA5.0.4^88^ were used for parameterisation and quantum mechanical (QM) calculations, respectively. Briefly, 40 GDP conformers were sampled in a 20 ns unrestrained NPT simulation at 450 K and 1 bar with the SPC/E_b_ water model^80^. The electrostatic potential in vacuum and explicit solvent (surrounding the solute within 5 Å) was then calculated for each conformation as previously described^40^ and partial charges fit to the electrostatic potentials with the mdgx programme^87^ while restraining the total charge to −3.0 and enforcing equal charges for chemically equivalent atoms. The vacuum and solvated partial charges were then averaged to obtain the IPolQ charges. The protocol was repeated with the resulting charges until the charges converge (< 10% change with respect to the previous iteration. The final partial charges used in this work are listed in Supplementary Table 1. We verified that these GDP parameters resulted in a stable HRAS-GDP complex for wild-type HRAS, including proper coordination of the bound Mg^2+^ ion, in agreement with the X-ray structure^50^. This was observed over microsecond-long simulations that are required for assessing fluorinated HRAS variants (**Fig. S4**).

### Long-timescale MD simulations of HemK

Starting from the crystal structure (PDB 1T43, residues 1-73), we centred the N-terminal domain (NTD) of HemK in a dodecahedral-shaped box at least 1.2 nm distanced from the edge of the box. Water molecules and MgCl_2_ (12mM) were then added to solvate and neutralise the system. The protein was parameterised with the DES-Amber force field^89^ and TIP4P-D water model^90^. Energy minimisation was performed using steepest descent. In the following MD runs, the leap-frog integrator and a 2 fs timestep along with the LINCS algorithm ^75^ to constrain all bonds connected to hydrogen were used. Nonbonded Van der Waals interactions were treated with 0.9 nm cut-off. Electrostatics were calculated using a 1.0 nm cut-off for the real-space contribution, and the Particle Mesh Ewald (PME) ^76^ method was used for long-range interactions with cubic interpolation and a fourierspacing of 0.125 nm. The temperature/pressure coupling parameters and equilibration protocol were as described above for the fluorinated protein variants. Six independent production simulations were launched starting with different initial velocities at the equilibration stage for 20 μs per replicate in the NPT ensemble (totalling 120 μs of sampling per variant). For the production runs, we employed a 4 fs timestep with hydrogen mass repartitioning applied^91^. Protein coordinates were saved and analysed every 1 ns, resulting in 20,000 snapshots per trajectory.

### Sidechain modelling of BTFMA

A model structure of a cysteine conjugated to BTFMA was first built in PyMol (version 2.3, Schrödinger, LLC) by mutating residue 655 in FLN5 (PDB 1QFH) to cysteine and completing the sidechain. The amino and carboxylate terminal groups were capped with acetyl and N-methyl amide groups respectively. ACPYPE was used to assign GAFF2 force field parameters^84,92^. Using the same nonbonded interaction cut-off values and equilibration protocols as for our AMBER-based MD simulations detailed above, we performed a 100 ns production simulation with position restraints (1000 kJ mol^-1^ nm^2^ along all coordinate axes) applied to the N, CA and C atoms to sample different sidechain conformations. Cys-BTFMA atoms were saved every 100 ps resulting in 1,000 snapshots of sidechain conformations. These were aligned to backbone atoms (N, CA, C) of residue 655 in FLN5, and conformations that resulted in clashes (< 2.5 Å for heavy atom distances) were discarded. Using these aligned rotamers we calculated the distribution of distances between the CF_3_ group of Cys-BTFMA and the Phe675 aromatic ring.

### AlphaFold structure predictions

We used the AlphaFold2^28^ and AlphaFold-Multimer^30^ implementation ColabFold v1.5.5^29^ to predict structures of all fluorinated protein variants and complexes by substituting tfmF with a tyrosine residue. All predictions were run on the Google Colab platform (https://colab.research.google.com/github/sokrypton/ColabFold/blob/main/batch/AlphaFold2_batch.ipy <u>nb</u>) using T4 GPUs and applying default settings with templates. All five models were relaxed for each variant. We also used AlphaFold3^31^ (https://alphafoldserver.com/) for all proteins and complexes. For HRAS, we explicitly included the ligands (Mg^2+^ and GDP). Default settings were applied.

### Analyses of MD simulations and predicted structures

We calculated the Cα-RMSD with respect to the energy minimised input structure for all simulations to ensure that the proteins remained stable and folded throughout the MD runs using the *gmx rms* tool^73^. The solvent-accessible surface area (SASA) for the CF_3_ group of tfmF was computed using the *gmx sasa* tool^73^. The root mean squared fluctuations (RMSF) of the protein backbone (Cα atoms) were computed using the gmx rmsf tool. The distances and angular orientation of the CF_3_ group (or tyrosine OH atom for AlphaFold predictions) relative to nearby aromatic sidechains were calculated using in-house python scripts and the MDAnalysis package^93^ (see Code Availability). We used the centre of mass of the CF_3_ group (or the tyrosine OH atom) and the centre of mass of the neighbouring aromatic ring heavy atoms for these calculations. Additionally, we calculated a geometric factor quantifying both the distance and angular contributions to the expected ring current effects within one term^51^, (1-3cos^2^θ)/r^3^, to more directly assess the correlation between the experimentally measured chemical shifts and computational predictions. Distances, angles and SASA values were calculated for every 1 ns in the trajectories. We calculated averages and standard errors from three independent simulation replicates, discarding the first 200 ns of each replicate to allow for equilibration of the sidechain rotamers. Positive variants were defined as variants that exhibited a secondary chemical shift of at least 0.2 ppm in magnitude. From the computational side, we defined positive predictions to have a distance of no more than 6 and 7 Å for perpendicular and in-plane interactions, respectively, with an orientational preference of more than 70% (i.e., at least 70% in-plane or 70% perpendicular). The distance cut-offs for AF2/AF3 predictions were 5.5 and 6.5 Å, respectively, to approximately account for the difference in non-clashing distances between a CF_3_ and OH group. Performance statistics were calculated based on these classifications.

## Data availability

All ^19^F NMR chemical shift data are available in the Supplementary Information. AlphaFold-generated structural models and MD trajectories are available on Zenodo at https://zenodo.org/records/14288915.

## Code availability

A Python script to calculate distances, angles and geometric factors for ring current predictions is available on GitHub (https://github.com/julian-streit/RingCurrents19F). MATLAB scripts to process and fit ^19^F NMR spectra are also available on GitHub (https://github.com/shschan/NMR-fit).

## Acknowledgements

This study was funded by a Wellcome Trust Investigator Award (to J.C., 206409/Z/17/Z). We acknowledge use of the UCL Biomolecular NMR Centre. Computational resources were provided by the Baskerville Tier 2 HPC service (https://www.baskerville.ac.uk/). Baskerville was funded by the EPSRC and UKRI through the World Class Labs scheme (EP/T022221/1) and the Digital Research Infrastructure programme (EP/W032244/1) and is operated by Advanced Research Computing at the University of Birmingham. We are also grateful to the UK Materials and Molecular Modelling Hub for computational resources, which is partially funded by the EPSRC (EP/T022213/1, EP/W032260/1 and EP/P020194/1), and the UCL Kathleen High Performance Computing Facility (Kathleen@UCL), and associated support services. We thank Prof. M. Rodnina (Max Planck Institute for Biophysical Chemistry, Göttingen) and Prof. D. Calderwood (Yale School of Medicine) for the kind gift of the HemK and human migfilin plasmids, respectively.

## Notes

https://zenodo.org/records/14288915

### Competing Interest Statement

The authors have declared no competing interest.

