## Supplementary Information for "Rational design of ^19^F NMR labelling sites to probe protein structure and interactions"

### Supplementary Figures

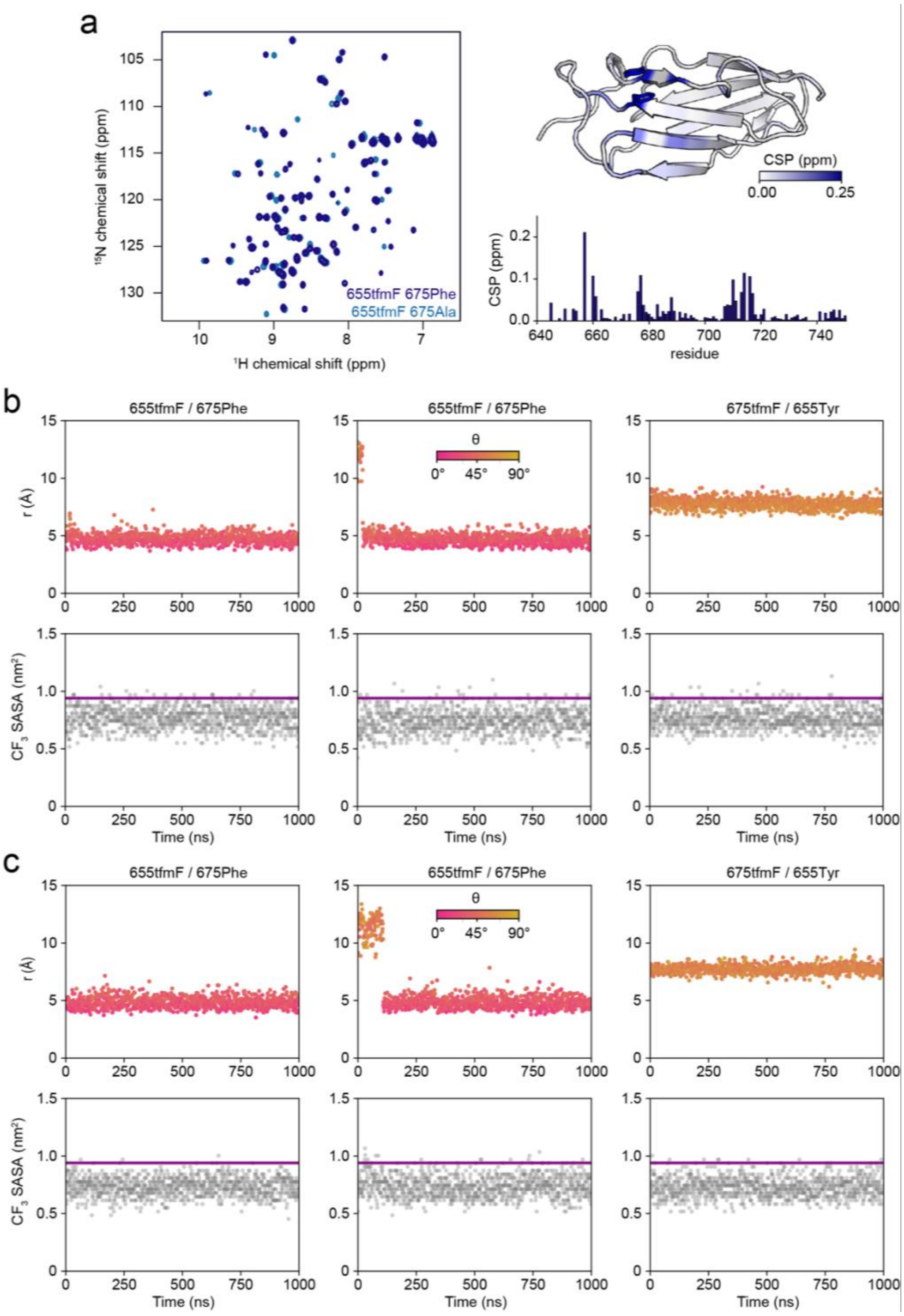


**Figure S1.** Experimental and computational characterisation of tfmF-labelled FLN5. **a.** 2D $^{1}$H,$^{15}$N SOFAST-HMQC spectra of FLN5 655tfmF with and without the F675A mutation, recorded at 298 K and 500 MHz. **b.** Distances (r) and angles (θ) between respective $CF_3$ groups and nearby aromatic rings observed in all-atom MD simulations with the ff15ipq and **c** C36m force field. The bottom panels show the solvent-accessible surface area (SASA) of the $CF_3$ group during the simulation and the horizontal line represents a fully solvated $CF_3$ group in a disordered peptide. The middle panels show simulation results of 655tfmF/675Phe initiated from a different sidechain rotamer of residue 655.

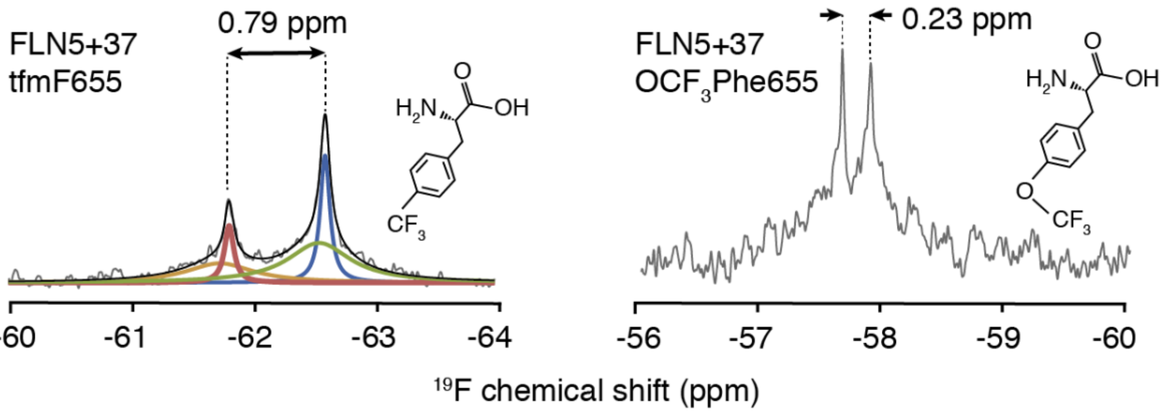


**Figure S2.** [19]F NMR of FLN5 ribosome-nascent chain complexes labelled with tfmF and tfmOF (trifluoromethoxy-L-phenylalanine) at residue 655. Spectra were recorded at 298 K and 500 MHz.

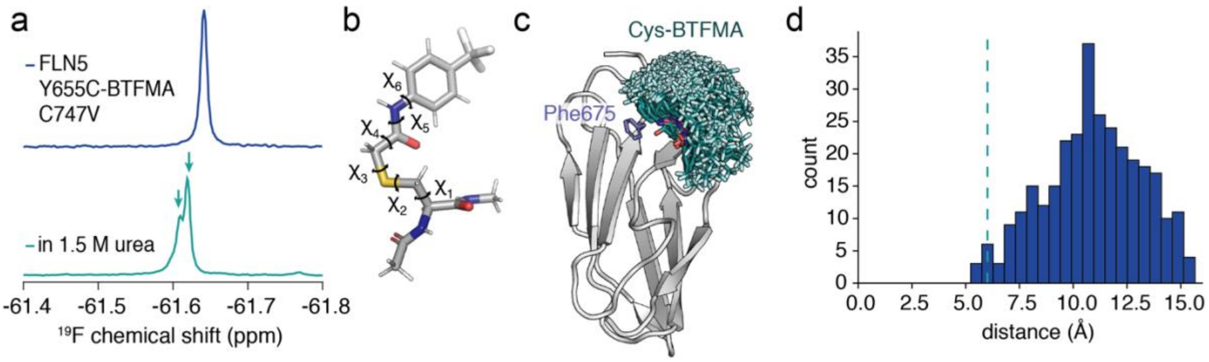


**Figure S3.** BTFMA-labelled FLN5. **a.** [19]F NMR spectrum of FLN5 655Cys labelled with 2-bromo-N-(4-[trifluoromethyl]phenyl)acetamide (BTFMA) (with a cysteine-free background, Cys747Val) in Tico buffer (top) and buffer with 1.5 M urea (bottom). The arrows highlight two separate peaks corresponding to the unfolded and folded conformation. **b.** Structure of a cysteine conjugated with BTFMA highlighting its rotatable sidechain dihedral angles. **c.** Cysteine-BTFMA rotamers aligned to residue 655 of the FLN5 crystal structure (PDB 1QFH) and **d.** a histogram of the $CF_3$-Phe675 ring distance (between the centres of mass). Dashed line indicates 6 Å, below which ring current effects are observable.

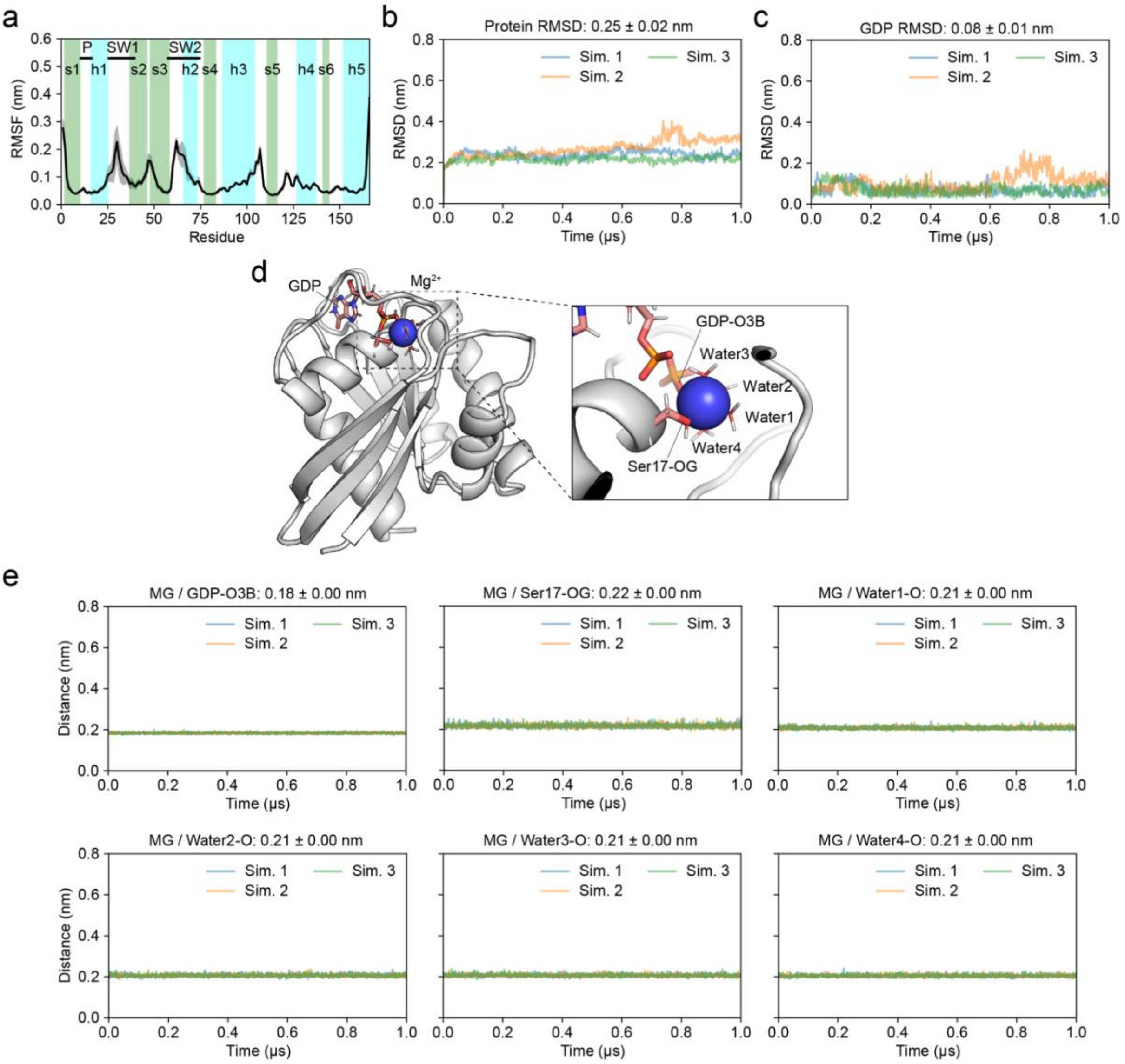


**Figure S4.** All-atom MD simulations of the HRAS-GDP complex with implicitly polarised charges (IPolQ) for GDP. **a.** Average root mean square fluctuations (RMSF) for each residue Cα atom (mean ± s.e.m. from three independent 1 μs-long simulations). **b-c.** Protein and ligand (all-atom) RMSD calculated for three independent 1 μs-long simulations (mean ± s.e.m.). **d.** Energy-minimised structure of HRAS-GDP highlighting the coordination of a structural $Mg^{2+}$ by a GDP phosphate oxygen atom, Ser17 oxygen atom and four structural water molecules. **e.** $Mg^{2+}$ coordination distances highlighting the stability of the structural ion and ligand in the complex (mean ± s.e.m.).

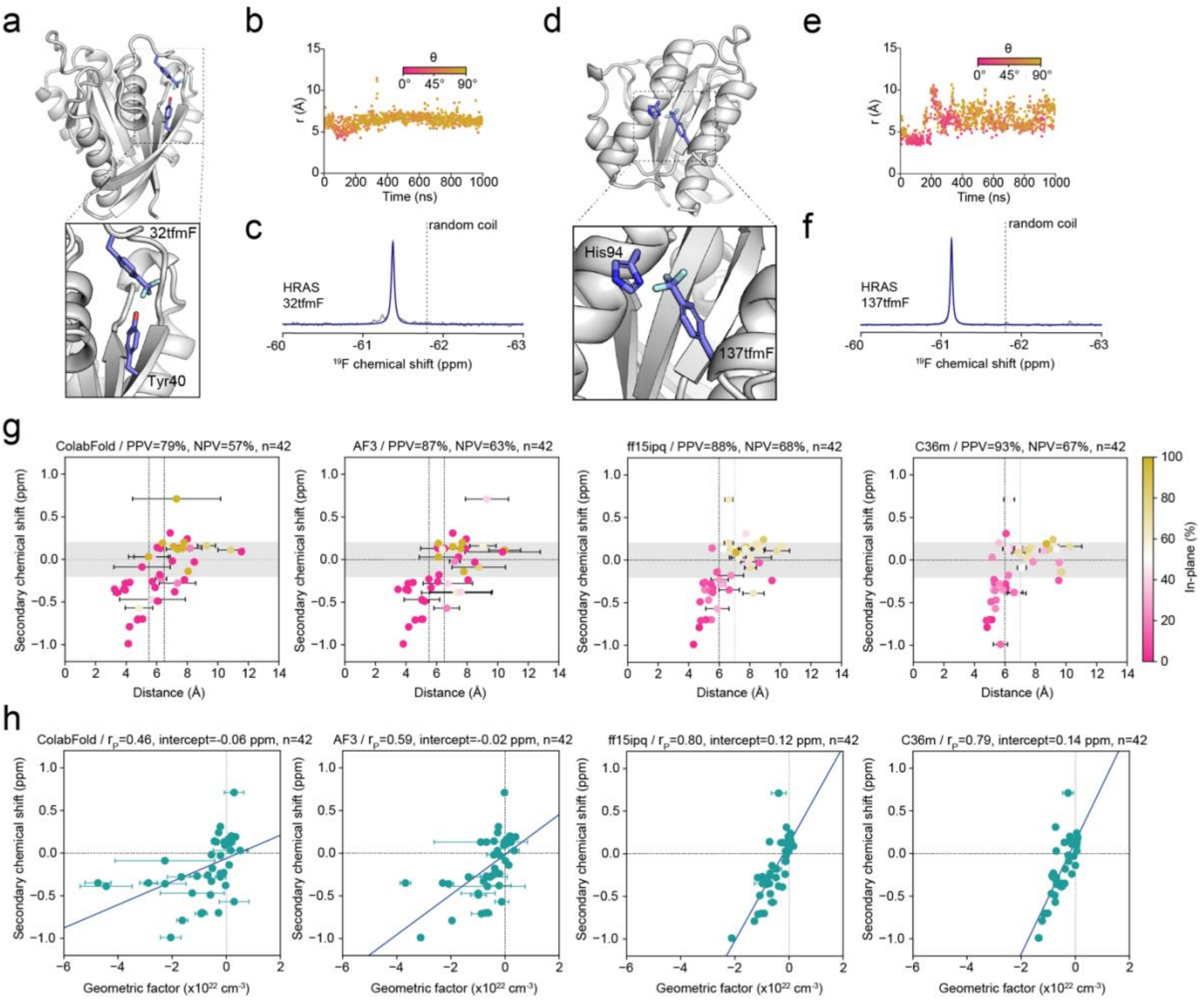


**Figure S5.** Ring current design to probe protein structure using $^{19}$F NMR chemical shifts. **a.** Structural model of HRAS 32tfmF, highlighting an interaction between tfmF and 40Tyr between a loop and β-strand. **b.** Distance (r) and angle (θ) between the $CF_3$ group of 32tfmF and aromatic ring of 40Tyr observed in a representative all-atom MD simulation. **c.** $^{19}$F NMR spectra of HRAS 32tfmF recorded at 298 K and 500 MHz. **d.** Structural model of HRAS 137tfmF highlighting an interaction between tfmF and 94His across two α-helices. **e.** Distance (r) and angle (θ) between the $CF_3$ group of 137tfmF and aromatic ring of 94His observed in a representative all-atom MD simulation. **f.** $^{19}$F NMR spectra of HRAS 137tfmF recorded at 298 K and 500 MHz. **g-h.** Correlation analysis for a subdataset of 42 proteins that could be modelled by both MD force fields (ff15ipq and C36m). **g.** Scatter plots correlating the distances (coloured by the fraction of time (MD) or models (ColabFold/AF3) spent in the plane of the ring defined as $\theta > 54.6°$) with secondary $^{19}$F chemical shifts for all protein variants predicted by ColabFold, AF3, ff15ipq (MD), and C36m (MD). The error bars represent one s.d. over the five predicted models for ColabFold and AF3, and the s.e.m. obtained from three independent simulations for MD. PPV=positive predictive value; NPV=negative predictive value. Positive secondary chemical shift > 0.2 ppm in magnitude. The vertical lines represent the distance cut-off values for perpendicular and in-plane interactions (lower and higher distance, respectively). **h.** Correlations between predicted geometric factors $(1\text{-}3\cos^2\theta)/r^3$) and secondary $^{19}$F chemical shifts and the corresponding Pearson correlation coefficients ($r_P$) and intercepts for lines of best fit.

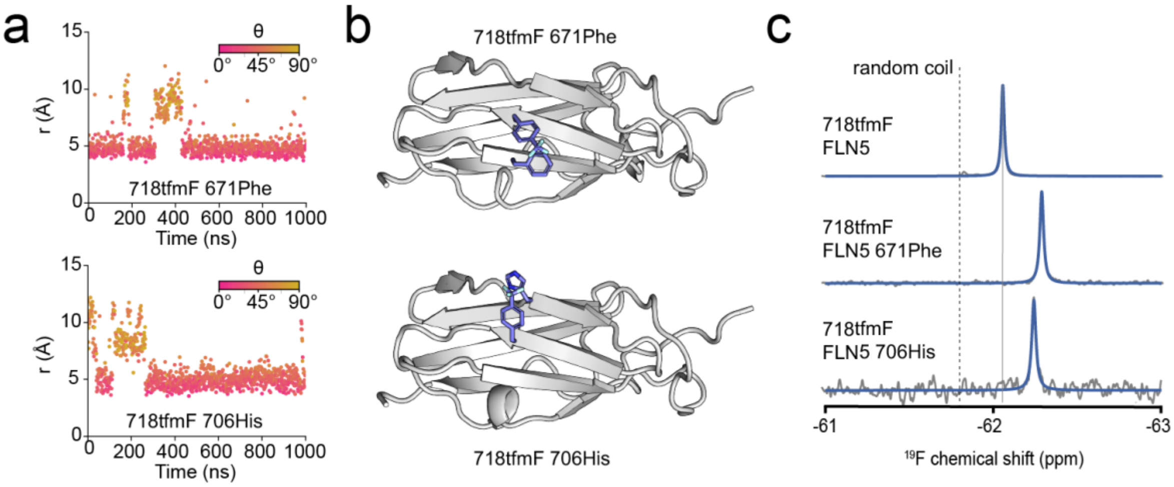


**Figure S6.** Probing different structural interactions using a single fluorine labelling site on FLN5. **a.** Distance (r) and angle (θ) between the $CF_3$ group of 718tfmF (E-strand) and aromatic ring of 671Phe (B-strand) and 706His (D-Strand) observed in representative all-atom MD simulations. **b.** Predicted structures of FLN5 718tfmF with 671Phe and 706His. **c.** $^{19}$F NMR spectra of FLN5 671tfmF on its own and with Glu671Phe and Lys706His point mutations, recorded at 298 K and 500 MHz.

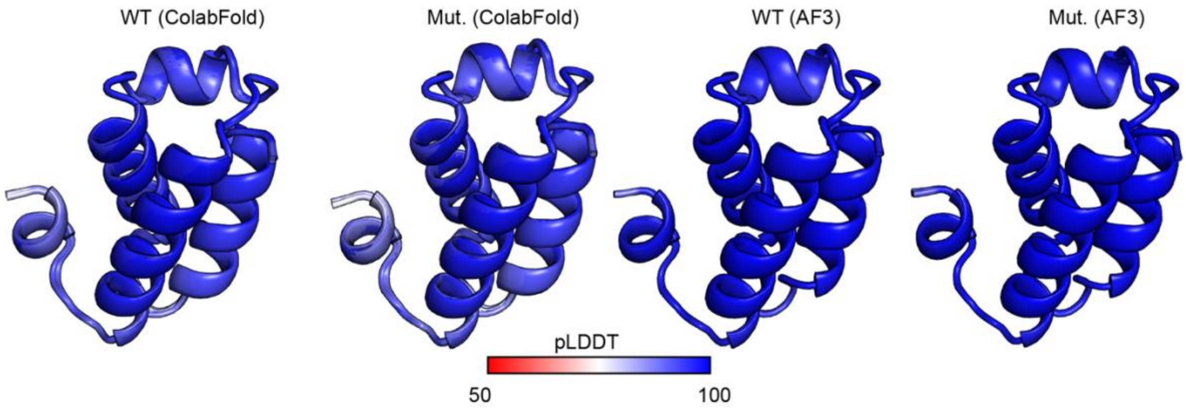


**Figure S7.** ColabFold and Alphafold3 (AF3) predictions of wild-type (WT) and mutant (Mut.) HemK coloured according to the pLDDT score. Default settings were used without templates for ColabFold.

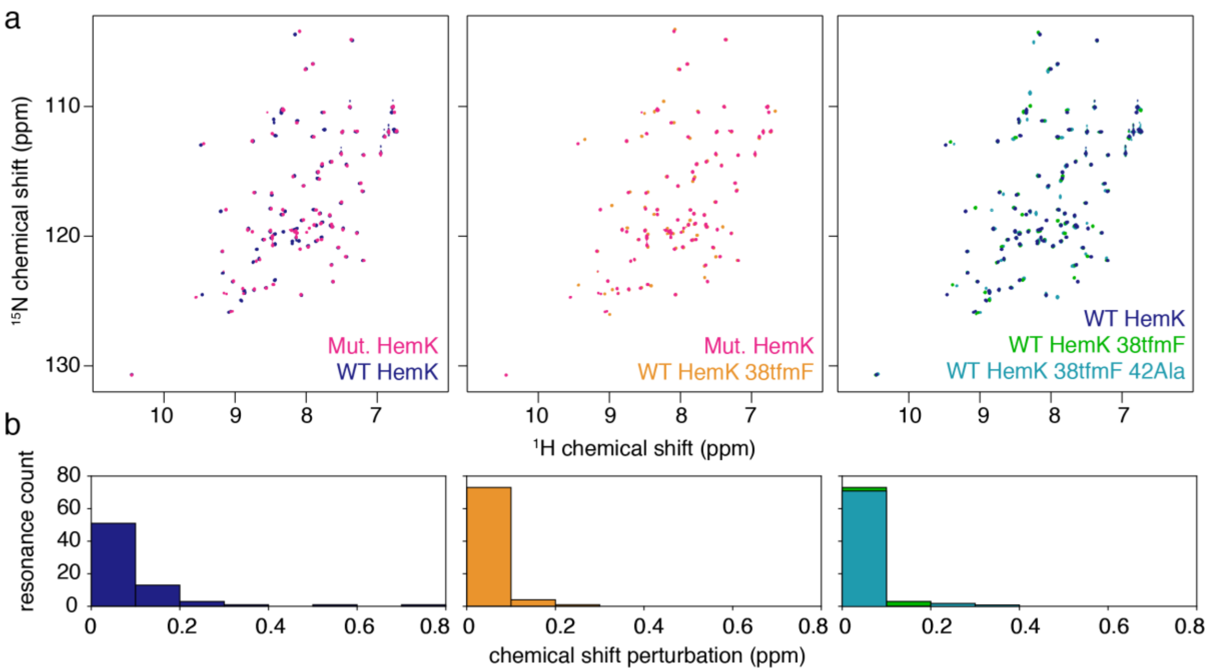


**Figure S8.** NMR characterisation of the HemK NTD. **a.** 2D $^{1}$H,$^{15}$N-HSQC spectra of HemK NTD variants, recorded at 298 K and 800 MHz. **b.** Histograms of chemical shift perturbations from analysis of spectra shown in a. Right histogram shows CSPs relative to wild-type (WT) HemK.

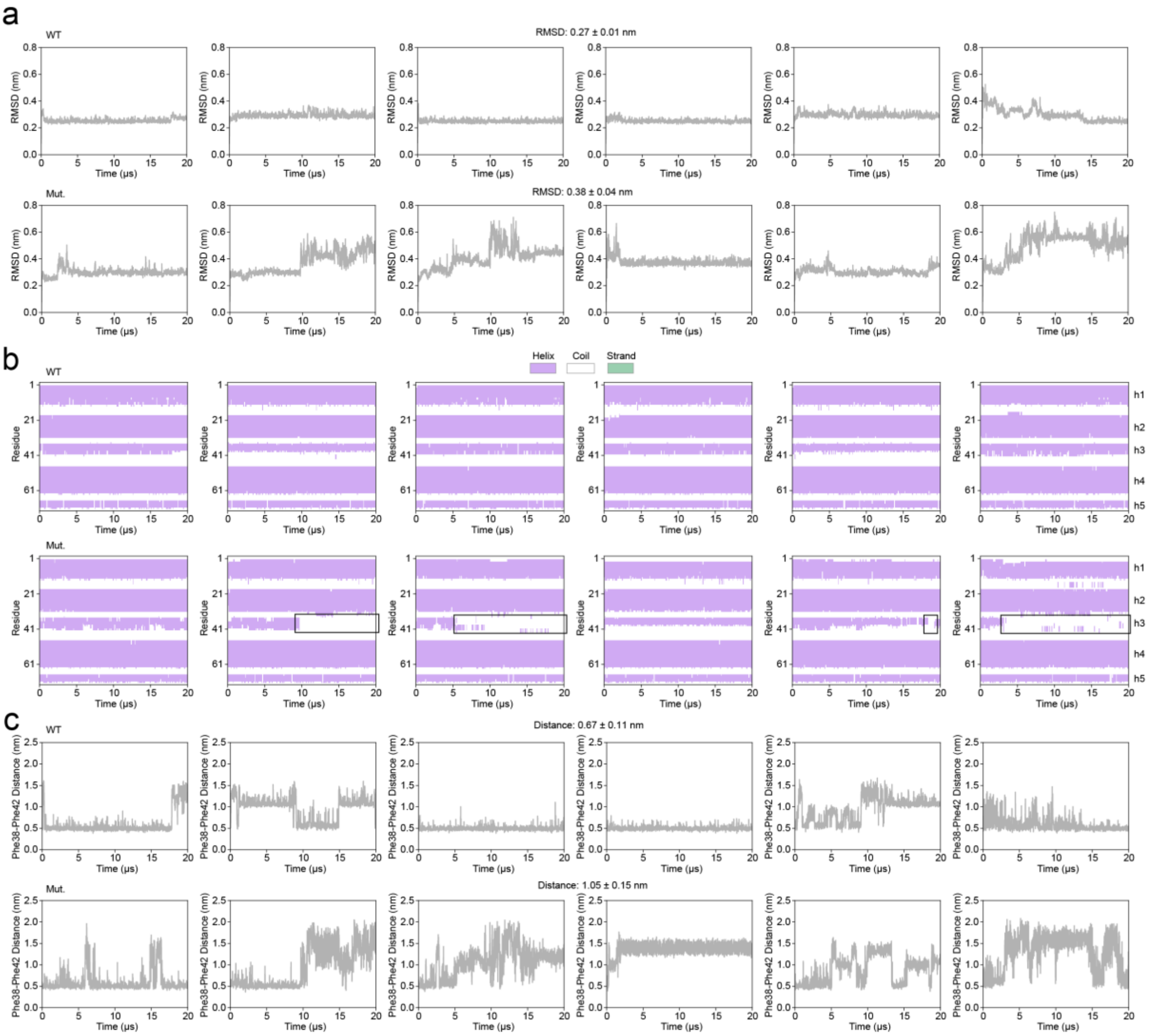


**Figure S9.** Long-timescale, all-atom MD simulations of HemK (residues 1-73, NTD). **a.** All-atom protein RMSD relative to the crystal structure (residues 1-73, PDB 1T43) of six independent MD simulations of 20 $\mu$s per variant. **b.** Secondary structure content (calculated with DSSP) of HemK observed during the MD simulations. The black boxes indicate parts of trajectories where helix h3 has completely unfolded for at least 1 $\mu$s. **c.** Distance between the centres of mass of the Phe38 and Phe42 aromatic rings. All averages in this figure represent the mean ± s.e.m.

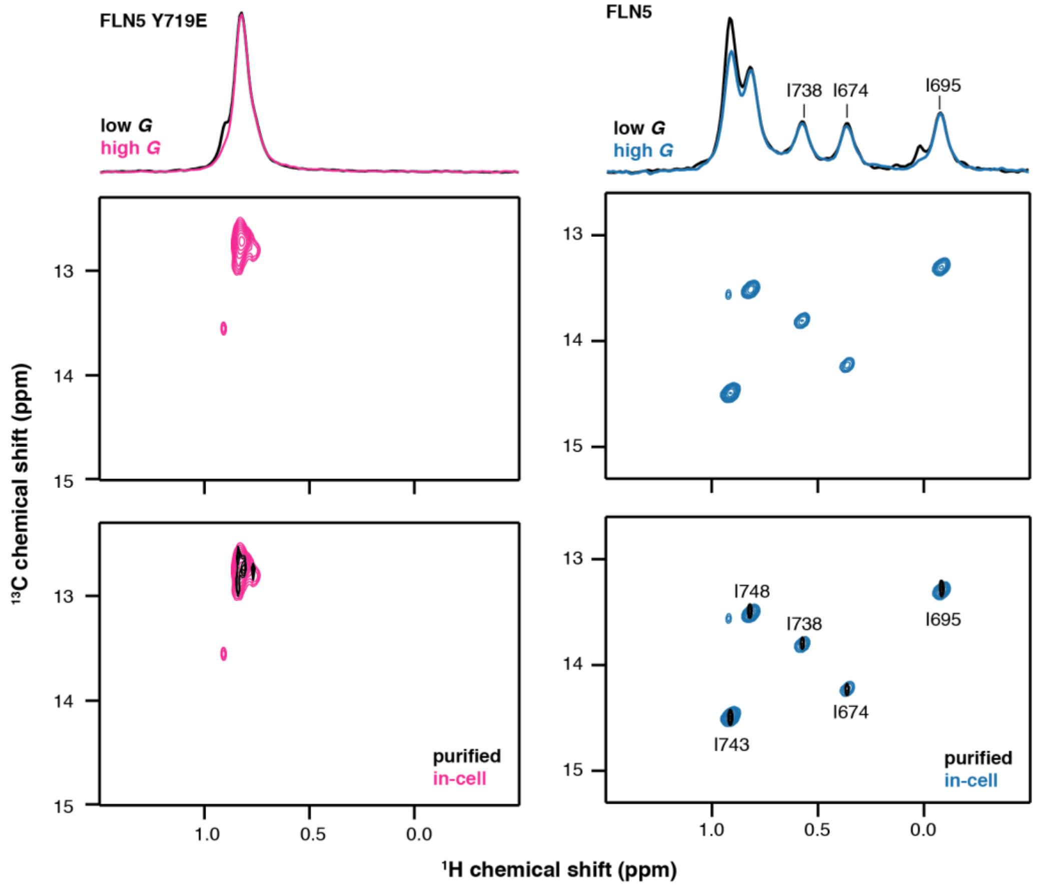


**Figure S10.** $^{1}H$,$^{13}C$ HMQC NMR spectra of uniform $^{2}H$, selectively Ileδ1-[$^{13}CH_3$]-labelled unfolded (left) FLN5 and folded (right) FLN5 recorded at 298K and 500 MHz. Top shows $^{1}H$,$^{13}C$ diffusion measurements to detect cell leakage (see Methods).

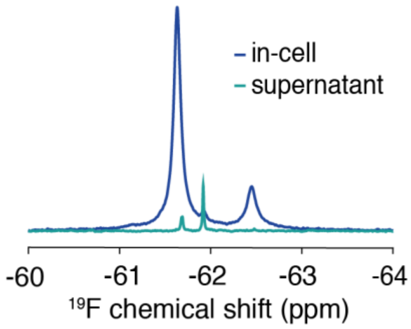


**Figure S11.** Quality control for in-cell NMR of FLN5 672A. Overlaid $^{19}F$ NMR spectra of in-cell FLN5 672A and the supernatant after centrifugation of the in-cell sample at the end of NMR acquisition recorded at 298 K and 500 MHz.

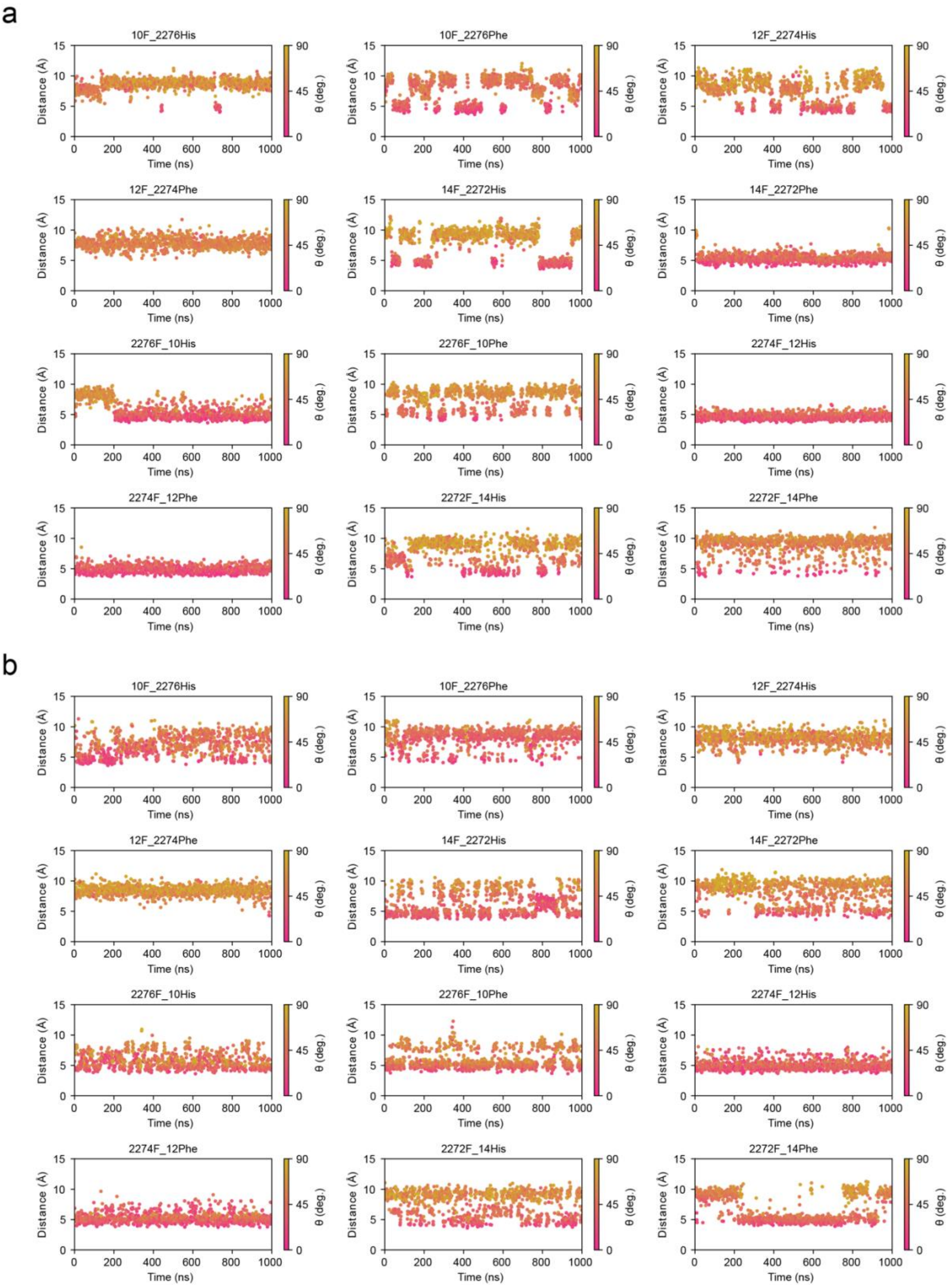


**Figure S12.** Using all-atom MD simulations of the FLNa21-migfilin complex to screen suitable fluorine labelling sites for protein-protein interaction detection with the **a.** ff15ipq and **b**. C36m force field.

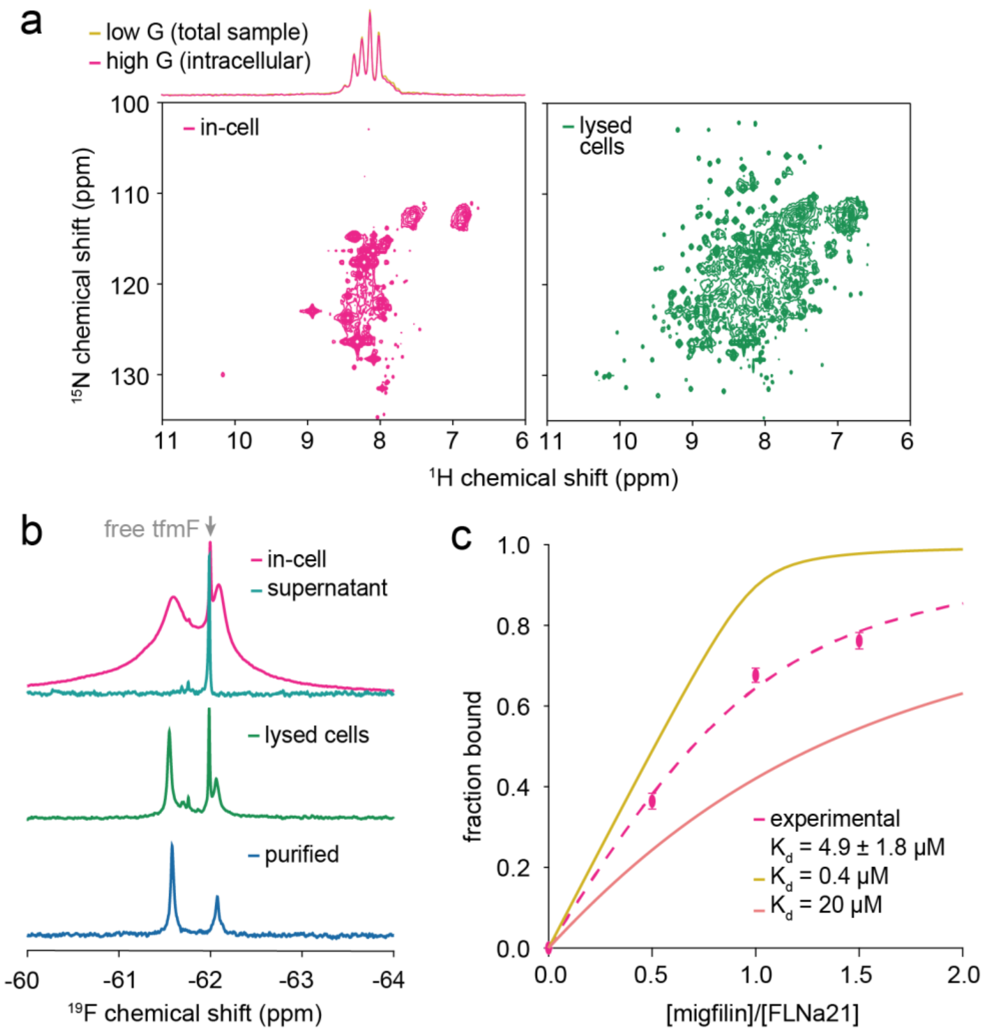


**Figure S13.** Characterisation of FLNa21-migfilin binding by NMR in living cells and buffer. **a.** In-cell and lysate 2D $^{1}H$,$^{15}N$ SOFAST-HMQC spectra of FLNa21 recorded at 298 K and 800 MHz. Top shows $^{1}H$,$^{15}N$-SORDID diffusion measurements to detect cell leakage (see Methods). **b.** $^{19}F$ NMR spectra of in-cell, lysate and purified FLNa21 2274tfmF co-expressed with migfilin 12His, including the supernatant obtained after centrifugation of the in-cell sample at the end of data acquisition. **c.** Titration and binding affinity fitting of migfilin 12His binding to FLNa21 2274tfmF (25 μM) under purified conditions (spectra shown in Figure 6e). Theoretical binding curves of an upper[94] and lower[47] bound binding affinities from the literature are shown as a comparison.

### Supplementary tables

| Atom name | GAFF2 atom type | Charge |
|---|---|---|
| PA | p5 | 1.856920 |
| PB | p5 | 2.238590 |
| C5’ | c3 | 0.133400 |
| O5’ | os | -0.607690 |
| C4’ | c3 | 0.260670 |
| O4’ | os | -0.514220 |
| C3’ | c3 | 0.269080 |
| O3’ | oh | -0.746930 |
| C2’ | c3 | 0.215010 |
| O2’ | oh | -0.746930 |
| C1’ | c3 | 0.269860 |
| N1 | ns | -0.763260 |
| O1A | o | -1.063290 |
| O1B | o | -1.237150 |
| C2 | cc | 0.896060 |
| N2 | nv | -0.950460 |
| O2A | o | -1.063290 |
| O2B | o | -1.237150 |
| N3 | nd | -0.770020 |
| O3A | os | -0.793990 |
| O3B | o | -1.237150 |
| C4 | cd | 0.397790 |
| C5 | cc | -0.018970 |
| C6 | c | 0.866040 |
| O6 | o | -0.785660 |
| N7 | nc | -0.713190 |
| C8 | cd | 0.300380 |
| N9 | na | -0.167520 |
| H1 | hn | 0.462020 |
| H8 | h5 | 0.155810 |
| H1’ | h2 | 0.098850 |
| H2’ | ho | 0.470330 |
| H21 | hn | 0.422320 |
| H3’ | h1 | 0.028270 |
| H22 | hn | 0.430200 |
| H4’ | h1 | 0.073600 |
| H5’ | h1 | 0.032150 |
| H3T | ho | 0.470330 |
| H2’’ | h1 | 0.037040 |
| H5’’ | h1 | 0.032150 |

**Table S1**. GDP partial charges used in this work (obtained with the IPolQ method).

| **Protein** | **Variant** | **$\Delta G_{folding}$ (kcal mol$^{-1}$)** |
|---|---|---|
| FLN5 | Wild-type | -7.01 ± 0.22 |
| FLN5 | 655F | -6.58 ± 0.25 |
| FLN5 | 673F 716His | -7.14 ± 0.28 |
| FLN5 | 675F | -7.22 ± 0.26 |
| FLN5 | 694F | -7.14 ± 0.26 |
| FLN5 | 718F 671Phe | -7.30 ± 0.25 |
| FLN5 | 718F 706His | -6.35 ± 0.25 |
| FLN5 | 726F 746Phe | -7.30 ± 0.25 |
| FLN5 | 728F 744His | -6.44 ± 0.26 |
| FLN5 | 732F | -5.81 ± 0.26 |
| HRAS (1-166) | Wild-type | -5.40 ± 0.10 |
| HRAS (1-166) | 32F | -4.58 ± 0.15 |
| HRAS (1-166) | 137F | -5.42 ± 0.17 |
| HRAS (1-166) | 157F 153His | -3.45 ± 0.11 |

**Table S2**. Folding free energies ($\Delta G_{folding}$) of fluorinated and wild-type proteins. The free energies were determined from the populations of unfolded and folded protein in urea relative to wild-type as measured by $^{19}$F NMR.

| Protein | Variant | Chemical shift (ppm) | Secondary shift (ppm) | r (Å) | r SD | % inplane | % inplane SD | tfmF pLDDT | Aromatic pLDDT | Geom. | Geom. SD |
|---|---|---|---|---|---|---|---|---|---|---|---|
| FLN5 | 655F | -62.61 | -0.79 | 4.22 | 0.1 | 0 | 0 | 98.82 | 98.74 | -1.62E+22 | 2.03E+21 |
| FLN5 | 665F_748Phe | -61.68 | 0.14 | 6.04 | 0.14 | 0 | 0 | 98.28 | 98.1 | -4.32E+21 | 3.81E+20 |
| FLN5 | 673F_716His | -62.17 | -0.35 | 3.84 | 0.08 | 0 | 0 | 98.84 | 98.84 | -2.88E+22 | 3.26E+21 |
| FLN5 | 673F_716Phe | -62.09 | -0.27 | 4.24 | 0.05 | 0 | 0 | 98.88 | 98.84 | -1.08E+22 | 2.69E+21 |
| FLN5 | 675F | -61.67 | 0.15 | 7.1 | 0.06 | 100 | 0 | 98.72 | 98.82 | 5.40E+20 | 1.43E+20 |
| FLN5 | 675F_714Phe | -61.69 | 0.13 | 6.24 | 0.1 | 0 | 0 | 98.58 | 98.22 | -3.70E+21 | 1.77E+21 |
| FLN5 | 692F_732Phe | -61.69 | 0.13 | 7.44 | 0.05 | 0 | 0 | 98.16 | 98.72 | -7.89E+20 | 6.02E+19 |
| FLN5 | 694F_730Phe | -61.69 | 0.13 | 8.18 | 0.77 | 20 | 40 | 98.5 | 98.6 | -6.47E+20 | 4.73E+20 |
| FLN5 | 694F_730His | -61.58 | 0.24 | 8 | 0.06 | 0 | 0 | 98.56 | 98.64 | -2.80E+21 | 2.48E+20 |
| FLN5 | 696F_728Phe | -61.96 | -0.14 | 8.06 | 0.14 | 100 | 0 | 98.52 | 98.3 | 1.08E+21 | 1.34E+20 |
| FLN5 | 710F_716His | -62.1 | -0.28 | 3.98 | 0.23 | 0 | 0 | 98.32 | 98.82 | -1.66E+22 | 5.14E+21 |
| FLN5 | 714F | -61.63 | 0.19 | 6.38 | 0.12 | 100 | 0 | 98.26 | 98.6 | 3.43E+21 | 1.17E+20 |
| FLN5 | 716F_673Phe | -61.51 | 0.31 | 6.88 | 0.1 | 0 | 0 | 98.84 | 98.88 | -2.17E+21 | 1.30E+20 |
| FLN5 | 718F_671Phe | -62.29 | -0.47 | 5.72 | 2.15 | 40 | 48.99 | 98.26 | 98.6 | -1.26E+22 | 1.18E+22 |
| FLN5 | 718F_706His | -62.21 | -0.39 | 3.4 | 0.24 | 0 | 0 | 97.96 | 98.3 | -4.44E+22 | 9.60E+21 |
| FLN5 | 718F_706Phe | -61.91 | -0.09 | 5.04 | 1.85 | 0 | 0 | 97.78 | 98.14 | -2.27E+22 | 1.84E+22 |
| FLN5 | 726F_746Phe | -62.52 | -0.7 | 5.08 | 0.04 | 0 | 0 | 98.02 | 97.9 | -2.85E+21 | 6.66E+20 |
| FLN5 | 728F_744His | -62.39 | -0.57 | 4.84 | 0.9 | 60 | 48.99 | 98.14 | 97.86 | 2.91E+21 | 5.46E+21 |
| FLN5 | 728F_744Phe | -61.79 | 0.03 | 5.7 | 1.19 | 40 | 48.99 | 98.04 | 97.6 | 1.71E+21 | 6.46E+21 |
| FLN5 | 732F_692Trp | -61.7 | 0.12 | 7.36 | 0.73 | 100 | 0 | 98.7 | 97.86 | 1.41E+21 | 3.35E+20 |
| FLN5 | 732F_692Phe | -61.69 | 0.13 | 7.68 | 0.1 | 100 | 0 | 98.76 | 98.4 | 2.09E+21 | 9.49E+19 |
| FLN5 | 732F_692His | -61.62 | 0.2 | 7.78 | 0.04 | 100 | 0 | 98.74 | 98.36 | 1.90E+21 | 5.13E+19 |
| FLN5 | 740F | -62.06 | -0.24 | 7.82 | 0.17 | 0 | 0 | 98.12 | 98.84 | -2.16E+21 | 1.97E+20 |
| FLN5 | 748F | -61.85 | -0.03 | 8.46 | 0.05 | 0 | 0 | 98.1 | 98.22 | -2.12E+21 | 2.47E+20 |
| FLN4 | 555F | -62.81 | -0.99 | 4.14 | 0.19 | 0 | 0 | 86.5 | 86.78 | -2.05E+22 | 3.76E+21 |
| FLN4 | 565F_646His | -62.05 | -0.23 | 6.28 | 0.36 | 20 | 40 | 86.38 | 89.38 | -9.47E+20 | 8.69E+20 |
| FLN4 | 616F_571His | -62.08 | -0.26 | 5.7 | 0.17 | 0 | 0 | 91.34 | 89.62 | -6.79E+21 | 8.47E+20 |
| FLN4 | 624F_644Phe | -62.52 | -0.7 | 4.78 | 0.1 | 0 | 0 | 92.8 | 91.62 | -8.87E+21 | 1.33E+21 |
| I27 | 5F_24Phe | -61.71 | 0.11 | 10.84 | 0.83 | 80 | 40 | 95.02 | 97.94 | -2.06E+20 | 8.66E+20 |
| I27 | 6F_24Phe | -61.84 | -0.02 | 7.02 | 0.07 | 0 | 0 | 97.94 | 98.28 | -5.47E+21 | 2.33E+20 |
| I27 | 9F_22Phe | -61.66 | 0.16 | 9.26 | 0.59 | 80 | 40 | 98.18 | 97.92 | 7.44E+20 | 7.22E+20 |
| I27 | 14F_87Phe | -62 | -0.18 | 6.2 | 0.13 | 0 | 0 | 98.38 | 98.14 | -5.09E+21 | 5.62E+20 |
| I27 | 14F_87His | -62.15 | -0.33 | 5.9 | 0.06 | 0 | 0 | 98.42 | 98.32 | -5.47E+21 | 5.26E+20 |
| I27 | 20F_61Trp | -61.73 | 0.09 | 11.54 | 0.08 | 0 | 0 | 98.7 | 98.58 | -4.55E+20 | 1.86E+19 |
| I27 | 59F | -62.18 | -0.36 | 3.94 | 0.14 | 0 | 0 | 98.7 | 98.5 | -2.26E+22 | 7.74E+21 |
| I27 | 72F_35His | -62.17 | -0.35 | 3.24 | 0.08 | 0 | 0 | 98.68 | 98.74 | -4.74E+22 | 4.77E+21 |
| FLNa21 | 2242F | -62.2 | -0.38 | 7.18 | 0.04 | 0 | 0 | 95.62 | 95.04 | -2.33E+20 | 1.02E+20 |
| FLNa21 | 2244F | -61.11 | 0.71 | 7.3 | 2.88 | 100 | 0 | 96.82 | 94.34 | 2.90E+21 | 3.57E+21 |
| FLNa21 | 2258F_2296Phe | -62.1 | -0.28 | 7.36 | 1.2 | 20 | 40 | 97.9 | 98.08 | -2.15E+21 | 1.88E+21 |
| FLNa21 | 2306F_2322His | -62.53 | -0.71 | 4.74 | 0.05 | 0 | 0 | 98.6 | 98.54 | -9.27E+21 | 2.23E+21 |
| HemK | 38F | -62.04 | -0.22 | 4.74 | 0.08 | 0 | 0 | 93.4 | 94.34 | -1.40E+22 | 1.49E+21 |
| HRAS | 5F_54His | -62.08 | -0.26 | 4.06 | 0.12 | 0 | 0 | 98 | 97.86 | -2.32E+22 | 3.34E+21 |
| HRAS | 32F | -61.37 | 0.45 | 13.56 | 2.98 | 100 | 0 | 66.26 | 93.28 | 5.49E+20 | 6.92E+20 |
| HRAS | 88F_92His | -62.28 | -0.46 | 3.64 | 0.08 | 20 | 40 | 96.46 | 94.34 | -1.48E+22 | 1.02E+22 |
| HRAS | 99F_103His | -61.73 | 0.09 | 4.1 | 0.17 | 20 | 40 | 91.44 | 91.4 | -7.31E+21 | 6.13E+21 |
| HRAS | 137F | -61.13 | 0.69 | 4.14 | 0.05 | 100 | 0 | 97.58 | 96.18 | 1.22E+22 | 2.18E+21 |
| HRAS | 157F_153His | -62.33 | -0.51 | 4.34 | 0.2 | 0 | 0 | 98.52 | 97.94 | -2.16E+22 | 3.57E+21 |
| FLNa21-Mig | 2274F | -61.55 | 0.03 | 5.46 | 1.3 | 100 | 0 | 97.2 | 94.46 | 5.13E+21 | 2.39E+21 |
| FLNa21-Mig | 2274F_12His | -62.07 | -0.49 | 6.06 | 0.23 | 0 | 0 | 97.2 | 95.34 | -5.83E+21 | 8.07E+20 |

**Table S3**. $^{19}$F NMR chemical shifts and descriptors calculated from ColabFold predictions. Geometric factors (Geom.) are given in cm$^{-3}$. HRAS 28F (28TAG) was not predicted because AF2/ColabFold does not model ligands.

| Protein | Variant | Chemical shift (ppm) | Secondary shift (ppm) | r (Å) | r SD | % inplane | % inplane SD | tfmF pLDDT | Aromatic pLDDT | Geom. | Geom. SD |
|---|---|---|---|---|---|---|---|---|---|---|---|
| FLN5 | 655F | -62.61 | -0.79 | 4.18 | 0.04 | 0 | 0 | 97.26 | 98.8 | -1.96E+22 | 7.03E+20 |
| FLN5 | 665F_748Phe | -61.68 | 0.14 | 6.06 | 0.05 | 0 | 0 | 95.4 | 97.66 | -4.08E+21 | 1.90E+20 |
| FLN5 | 673F_716His | -62.17 | -0.35 | 4 | 0 | 0 | 0 | 94.84 | 97.8 | -2.31E+22 | 8.49E+20 |
| FLN5 | 673F_716Phe | -62.09 | -0.27 | 4.46 | 0.08 | 0 | 0 | 96.28 | 98.5 | -6.85E+21 | 2.30E+21 |
| FLN5 | 675F | -61.67 | 0.15 | 7.06 | 0.05 | 100 | 0 | 97.94 | 98.42 | 5.81E+20 | 4.19E+19 |
| FLN5 | 675F_714Phe | -61.69 | 0.13 | 6.12 | 0.04 | 0 | 0 | 96.58 | 96.02 | -6.88E+21 | 6.46E+20 |
| FLN5 | 692F_732Phe | -61.69 | 0.13 | 6.52 | 1.56 | 0 | 0 | 69.16 | 98.58 | -8.96E+21 | 1.72E+22 |
| FLN5 | 694F_730Phe | -61.69 | 0.13 | 7.9 | 0 | 0 | 0 | 91.42 | 93.9 | -2.39E+20 | 6.41E+19 |
| FLN5 | 694F_730His | -61.58 | 0.24 | 7.94 | 0.05 | 0 | 0 | 87.94 | 93.12 | -2.69E+21 | 7.02E+19 |
| FLN5 | 696F_728Phe | -61.96 | -0.14 | 7.78 | 0.04 | 100 | 0 | 88.94 | 89.4 | 1.37E+21 | 4.33E+19 |
| FLN5 | 710F_716His | -62.1 | -0.28 | 4.14 | 0.05 | 0 | 0 | 92.78 | 95.24 | -1.35E+22 | 3.78E+20 |
| FLN5 | 714F | -61.63 | 0.19 | 6.12 | 0.04 | 100 | 0 | 78.48 | 98.44 | 3.88E+21 | 2.16E+20 |
| FLN5 | 716F_673Phe | -61.51 | 0.31 | 7.06 | 0.08 | 0 | 0 | 95.16 | 98.4 | -2.44E+21 | 9.53E+19 |
| FLN5 | 718F_671Phe | -62.29 | -0.47 | 5.04 | 1.16 | 0 | 0 | 56.1 | 89.18 | -9.92E+21 | 6.28E+21 |
| FLN5 | 718F_706His | -62.21 | -0.39 | 7.3 | 2.28 | 60 | 48.99 | 53.3 | 87.52 | -6.52E+21 | 1.40E+22 |
| FLN5 | 718F_706Phe | -61.91 | -0.09 | 8.76 | 1.75 | 80 | 40 | 54.16 | 89.84 | 1.49E+20 | 1.08E+20 |
| FLN5 | 726F_746Phe | -62.52 | -0.7 | 5 | 0 | 0 | 0 | 90.94 | 96.72 | -7.16E+21 | 5.91E+20 |
| FLN5 | 728F_744His | -62.39 | -0.57 | 6.66 | 0.83 | 20 | 40 | 61.52 | 89.74 | -1.22E+21 | 2.53E+21 |
| FLN5 | 728F_744Phe | -61.79 | 0.03 | 7.44 | 0.14 | 0 | 0 | 81.6 | 93.64 | -2.80E+21 | 8.95E+19 |
| FLN5 | 732F_692Trp | -61.7 | 0.12 | 6.38 | 0.26 | 60 | 48.99 | 96.32 | 89.32 | 4.14E+20 | 7.23E+20 |
| FLN5 | 732F_692Phe | -61.69 | 0.13 | 7.58 | 0.04 | 100 | 0 | 96.34 | 96.5 | 2.22E+21 | 3.59E+19 |
| FLN5 | 732F_692His | -61.62 | 0.2 | 7.68 | 0.04 | 100 | 0 | 95.72 | 92.12 | 1.82E+21 | 7.01E+19 |
| FLN5 | 740F | -62.06 | -0.24 | 8.06 | 0.08 | 0 | 0 | 86.9 | 98.48 | -2.02E+21 | 8.65E+19 |
| FLN5 | 748F | -61.85 | -0.03 | 8.5 | 0 | 0 | 0 | 94.5 | 97.9 | -1.72E+21 | 3.58E+19 |
| FLN4 | 555F | -62.81 | -0.99 | 3.82 | 0.04 | 0 | 0 | 97.24 | 98.54 | -3.12E+22 | 1.04E+21 |
| FLN4 | 565F_646His | -62.05 | -0.23 | 5.5 | 0.09 | 0 | 0 | 93.98 | 97.14 | -3.39E+21 | 4.31E+20 |
| FLN4 | 616F_571His | -62.08 | -0.26 | 6.08 | 0.04 | 0 | 0 | 95.92 | 91.56 | -4.63E+21 | 1.43E+20 |
| FLN4 | 624F_644Phe | -62.52 | -0.7 | 5.1 | 0 | 0 | 0 | 94.64 | 97.98 | -6.30E+21 | 2.09E+20 |
| I27 | 5F_24Phe | -61.71 | 0.11 | 10.46 | 1.04 | 100 | 0 | 70.74 | 89.96 | 7.23E+20 | 4.68E+20 |
| I27 | 6F_24Phe | -61.84 | -0.02 | 7.2 | 0.35 | 20 | 40 | 63.9 | 94.16 | -4.14E+21 | 2.59E+21 |
| I27 | 9F_22Phe | -61.66 | 0.16 | 9.04 | 0.85 | 60 | 48.99 | 80.5 | 87.38 | 4.73E+20 | 7.50E+20 |
| I27 | 14F_87Phe | -62 | -0.18 | 6.16 | 0.05 | 0 | 0 | 93.34 | 96.92 | -3.79E+21 | 1.85E+20 |
| I27 | 14F_87His | -62.15 | -0.33 | 5.64 | 0.1 | 0 | 0 | 93.28 | 96.76 | -7.87E+21 | 1.18E+21 |
| I27 | 20F_61Trp | -61.73 | 0.09 | 10.32 | 2.46 | 0 | 0 | 75.62 | 96.46 | -4.29E+20 | 8.30E+19 |
| I27 | 59F | -62.18 | -0.36 | 4.14 | 0.05 | 0 | 0 | 95.54 | 96.06 | -2.08E+22 | 1.87E+21 |
| I27 | 72F_35His | -62.17 | -0.35 | 3.48 | 0.04 | 0 | 0 | 91.88 | 97.72 | -3.68E+22 | 1.94E+21 |
| FLNa21 | 2242F | -62.2 | -0.38 | 7.6 | 2.05 | 40 | 48.99 | 56.94 | 93.16 | -2.17E+21 | 4.43E+21 |
| FLNa21 | 2244F | -61.11 | 0.71 | 9.3 | 1.41 | 40 | 48.99 | 63.8 | 90.74 | -2.01E+20 | 6.51E+20 |
| FLNa21 | 2258F_2296Phe | -62.1 | -0.28 | 6.74 | 1.61 | 40 | 48.99 | 57.38 | 92.32 | -5.50E+21 | 6.28E+21 |
| FLNa21 | 2306F_2322His | -62.53 | -0.71 | 4.6 | 0.18 | 0 | 0 | 91.58 | 96.84 | -8.70E+21 | 3.74E+21 |
| HemK | 38F | -62.04 | -0.22 | 5 | 0 | 0 | 0 | 95.06 | 96.98 | -9.82E+21 | 1.47E+20 |
| HRAS | 5F_54His | -62.08 | -0.26 | 3.9 | 0.13 | 0 | 0 | 89.82 | 95.52 | -2.62E+22 | 2.74E+21 |
| HRAS | 28F | -62.24 | -0.46 | 3.54 | 0.05 | 0 | 0 | 93.94 | 98.36 | -2.33E+22 | 1.70E+21 |
| HRAS | 32F | -61.37 | 0.45 | 7.52 | 3.89 | 100 | 0 | 51.96 | 93.98 | 4.65E+21 | 2.26E+21 |
| HRAS | 88F_92His | -62.28 | -0.46 | 4.06 | 0.37 | 0 | 0 | 84.76 | 88.08 | -1.84E+22 | 1.31E+22 |
| HRAS | 99F_103His | -61.73 | 0.09 | 3.88 | 0.04 | 0 | 0 | 72.1 | 86.92 | -1.60E+22 | 3.20E+21 |
| HRAS | 137F | -61.13 | 0.69 | 4.36 | 0.87 | 100 | 0 | 95.08 | 91.38 | 1.40E+22 | 4.89E+21 |
| HRAS | 157F_153His | -62.33 | -0.51 | 4.84 | 0.15 | 100 | 0 | 87.36 | 89.76 | 8.67E+21 | 7.44E+20 |
| FLNa21-Mig | 2274F | -61.55 | 0.03 | 6.12 | 1.25 | 100 | 0 | 88.72 | 91.98 | 3.36E+21 | 1.86E+21 |
| FLNa21-Mig | 2274F_12His | -62.07 | -0.49 | 5.22 | 0.29 | 0 | 0 | 89.5 | 94.24 | -9.85E+21 | 2.85E+21 |

**Table S4**. $^{19}$F NMR chemical shifts and descriptors calculated from AF3 predictions. Geometric factors (Geom.) are given in cm$^{-3}$.

| Protein | Variant | Chemical shift (ppm) | Secondary shift (ppm) | r (Å) | r SEM | % inplane | % inplane SEM | Geom. | Geom. SEM |
|---|---|---|---|---|---|---|---|---|---|
| FLN5 | 655F | -62.61 | -0.79 | 4.69 | 0.02 | 0.2 | 0.12 | -1.28E+22 | 3.23E+20 |
| FLN5 | 665F_748Phe | -61.68 | 0.14 | 5.53 | 0.01 | 13.23 | 0.78 | -7.18E+21 | 2.03E+20 |
| FLN5 | 673F_716His | -62.17 | -0.35 | 5.06 | 0.02 | 18.07 | 0.9 | -7.25E+21 | 3.79E+20 |
| FLN5 | 673F_716Phe | -62.09 | -0.27 | 4.95 | 0.01 | 5.1 | 0.57 | -9.83E+21 | 4.78E+20 |
| FLN5 | 675F | -61.67 | 0.15 | 7.73 | 0.02 | 81.77 | 0.67 | 6.60E+20 | 9.62E+18 |
| FLN5 | 675F_714Phe | -61.69 | 0.13 | 6.93 | 0.53 | 48.07 | 20.67 | -2.23E+21 | 1.59E+21 |
| FLN5 | 692F_732Phe | -61.69 | 0.13 | 8.6 | 0.4 | 55.57 | 4.37 | -7.22E+20 | 3.14E+20 |
| FLN5 | 694F_730Phe | -61.69 | 0.13 | 7.7 | 0.4 | 53.17 | 8.95 | -2.76E+21 | 1.39E+21 |
| FLN5 | 694F_730His | -61.58 | 0.24 | 8.89 | 0.13 | 73.67 | 8.61 | 5.39E+20 | 1.78E+20 |
| FLN5 | 696F_728Phe | -61.96 | -0.14 | 5.85 | 0.18 | 9.5 | 4.14 | -6.23E+21 | 3.73E+20 |
| FLN5 | 710F_716His | -62.1 | -0.28 | 6.17 | 0.23 | 28.7 | 3.63 | -4.78E+21 | 9.01E+20 |
| FLN5 | 714F | -61.63 | 0.19 | 8.42 | 0.15 | 57.93 | 11.94 | 3.93E+20 | 3.12E+20 |
| FLN5 | 716F_673Phe | -61.51 | 0.31 | 7.74 | 0.16 | 36.43 | 4.76 | -9.50E+20 | 2.49E+20 |
| FLN5 | 718F_671Phe | -62.29 | -0.47 | 5.3 | 0.12 | 17 | 3.49 | -6.90E+21 | 5.75E+20 |
| FLN5 | 718F_706His | -62.21 | -0.39 | 8.24 | 0.69 | 57.23 | 4.88 | -2.49E+21 | 1.24E+21 |
| FLN5 | 718F_706Phe | -61.91 | -0.09 | 8.04 | 0.39 | 62.03 | 8.25 | -1.35E+21 | 1.34E+21 |
| FLN5 | 726F_746Phe | -62.52 | -0.7 | 5.48 | 0.19 | 14.6 | 1.91 | -9.14E+21 | 6.70E+20 |
| FLN5 | 728F_744His | -62.39 | -0.57 | 5.87 | 0.76 | 26.7 | 8.49 | -6.13E+21 | 1.54E+21 |
| FLN5 | 728F_744Phe | -61.79 | 0.03 | 7.28 | 0.47 | 51.7 | 4.36 | -1.29E+21 | 7.51E+20 |
| FLN5 | 732F_692Trp | -61.7 | 0.12 | 6.8 | 0.39 | 59.57 | 7.24 | -1.44E+20 | 2.68E+20 |
| FLN5 | 732F_692Phe | -61.69 | 0.13 | 6.29 | 0.09 | 43.67 | 1.9 | -9.36E+20 | 7.66E+19 |
| FLN5 | 732F_692His | -61.62 | 0.2 | 6.62 | 0.29 | 59.37 | 4.1 | -2.46E+20 | 1.51E+20 |
| FLN5 | 740F | -62.06 | -0.24 | 9.45 | 0.12 | 4.4 | 0.61 | -1.43E+21 | 8.51E+19 |
| FLN5 | 748F | -61.85 | -0.03 | 8.62 | 0.15 | 9.9 | 0.31 | -1.17E+21 | 8.62E+18 |
| FLN4 | 555F | -62.81 | -0.99 | 4.31 | 0.03 | 0.03 | 0.03 | -2.11E+22 | 8.62E+20 |
| FLN4 | 565F_646His | -62.05 | -0.23 | 6.02 | 0.57 | 22.4 | 6.37 | -6.98E+21 | 8.74E+20 |
| FLN4 | 616F_571His | -62.08 | -0.26 | 5.07 | 0.01 | 8.17 | 0.33 | -7.20E+21 | 1.74E+20 |
| FLN4 | 624F_644Phe | -62.52 | -0.7 | 5.12 | 0.02 | 3.87 | 0.24 | -1.02E+22 | 1.27E+20 |
| I27 | 5F_24Phe | -61.71 | 0.11 | 9.77 | 0.83 | 63.57 | 14.92 | -5.41E+20 | 4.61E+20 |
| I27 | 6F_24Phe | -61.84 | -0.02 | 7.9 | 0.49 | 35.87 | 7.83 | -1.04E+21 | 4.71E+20 |
| I27 | 9F_22Phe | -61.66 | 0.16 | 10.1 | 0.14 | 58.83 | 5.41 | 2.54E+20 | 5.93E+19 |
| I27 | 14F_87Phe | -62 | -0.18 | 6.77 | 0.82 | 19.7 | 11.36 | -4.58E+21 | 1.81E+21 |
| I27 | 14F_87His | -62.15 | -0.33 | 5.58 | 0.02 | 6.07 | 0.47 | -8.55E+21 | 3.69E+19 |
| I27 | 20F_61Trp | -61.73 | 0.09 | 7.1 | 0.04 | 87.7 | 2.28 | 1.66E+21 | 1.21E+20 |
| I27 | 59F | -62.18 | -0.36 | 5.4 | 0.23 | 12.23 | 2.4 | -8.52E+21 | 9.73E+20 |
| I27 | 72F_35His | -62.17 | -0.35 | 6.65 | 0.67 | 9.9 | 2.26 | -9.77E+21 | 3.02E+21 |
| FLNa21 | 2242F | -62.2 | -0.38 | 5.57 | 0.03 | 7.83 | 0.91 | -3.84E+21 | 3.38E+19 |
| FLNa21 | 2244F | -61.11 | 0.71 | 6.63 | 0.26 | 55.23 | 12.43 | -3.83E+21 | 2.73E+21 |
| FLNa21 | 2258F_2296Phe | -62.1 | -0.28 | 5.47 | 0.15 | 5.7 | 2.33 | -1.16E+22 | 5.01E+20 |
| FLNa21 | 2306F_2322His | -62.53 | -0.71 | 4.79 | 0.02 | 5.27 | 0.15 | -1.14E+22 | 1.54E+20 |
| HemK | 38F | -62.04 | -0.22 | 5.14 | 0.02 | 1.13 | 0.44 | -9.79E+21 | 4.77E+20 |
| HRAS | 5F_54His | -62.08 | -0.26 | 4.56 | 0.02 | 0.8 | 0.12 | -1.41E+22 | 4.24E+20 |
| HRAS | 28F | -62.24 | -0.46 | 4.41 | 0.07 | 0.13 | 0.11 | -1.94E+22 | 2.40E+20 |
| HRAS | 32F | -61.37 | 0.45 | 6.32 | 0.45 | 83.57 | 11.47 | 7.06E+20 | 1.65E+21 |
| HRAS | 88F_92His | -62.28 | -0.46 | 11.25 | 0.11 | 89.33 | 1.47 | 5.07E+20 | 1.16E+20 |
| HRAS | 99F_103His | -61.73 | 0.09 | 6.17 | 0.33 | 28.53 | 3.65 | -2.89E+21 | 5.00E+20 |
| HRAS | 137F | -61.13 | 0.69 | 6.73 | 0.17 | 66.53 | 4.45 | -4.68E+20 | 9.00E+20 |
| HRAS | 157F_153His | -62.33 | -0.51 | 5.4 | 0.09 | 53.1 | 9.44 | -1.98E+21 | 1.33E+21 |
| FLNa21-Mig | 2274F | -61.55 | 0.03 | 8.04 | 1.37 | 56.37 | 3.7 | -4.36E+19 | 1.24E+20 |
| FLNa21-Mig | 2274F_12His | -62.07 | -0.49 | 4.75 | 0.01 | 6.1 | 0.79 | -1.08E+22 | 2.66E+19 |

**Table S5.** $^{19}$F NMR chemical shifts and descriptors calculated from MD (ff15ipq) predictions. Geometric factors (Geom.) are given in cm$^{-3}$.

| Protein | Variant | Chemical shift (ppm) | Secondary shift (ppm) | r (Å) | r SEM | % inplane | % inplane SEM | Geom. | Geom. SEM |
|---|---|---|---|---|---|---|---|---|---|
| FLN5 | 655F | -62.61 | -0.79 | 4.83 | 0.01 | 0.03 | 0.03 | -1.22E+22 | 1.63E+20 |
| FLN5 | 665F_748Phe | -61.68 | 0.14 | 6.89 | 0.24 | 40.27 | 2.92 | -2.17E+21 | 2.94E+20 |
| FLN5 | 673F_716His | -62.17 | -0.35 | 5.21 | 0.05 | 33.3 | 1.12 | -3.91E+21 | 1.57E+20 |
| FLN5 | 673F_716Phe | -62.09 | -0.27 | 5.31 | 0.17 | 9.7 | 2.92 | -8.39E+21 | 4.82E+20 |
| FLN5 | 675F | -61.67 | 0.15 | 7.7 | 0.01 | 66.27 | 2.38 | 2.89E+20 | 5.63E+19 |
| FLN5 | 675F_714Phe | -61.69 | 0.13 | 6.31 | 0.05 | 23.47 | 9.08 | -4.04E+21 | 9.72E+20 |
| FLN5 | 692F_732Phe | -61.69 | 0.13 | 9.38 | 0.09 | 45.03 | 0.81 | -1.28E+20 | 1.71E+19 |
| FLN5 | 694F_730Phe | -61.69 | 0.13 | 8.9 | 0.03 | 76 | 0.37 | 5.83E+20 | 2.32E+19 |
| FLN5 | 694F_730His | -61.58 | 0.24 | 9.14 | 0.02 | 77.83 | 0.82 | 5.33E+20 | 1.39E+19 |
| FLN5 | 696F_728Phe | -61.96 | -0.14 | 9.68 | 0.19 | 72.83 | 3.1 | 7.05E+18 | 2.07E+20 |
| FLN5 | 710F_716His | -62.1 | -0.28 | 6.04 | 0.07 | 6.23 | 0.81 | -6.96E+21 | 1.68E+20 |
| FLN5 | 714F | -61.63 | 0.19 | 8.71 | 0.02 | 93.93 | 0.78 | 8.90E+20 | 1.56E+19 |
| FLN5 | 716F_673Phe | -61.51 | 0.31 | 6.07 | 0.14 | 5.97 | 1.79 | -7.31E+21 | 7.39E+20 |
| FLN5 | 718F_671Phe | -62.29 | -0.47 | 5.41 | 0.18 | 15.73 | 3.35 | -8.85E+21 | 5.80E+20 |
| FLN5 | 718F_706His | -62.21 | -0.39 | 6.76 | 0.61 | 38.27 | 9.87 | -5.12E+21 | 1.41E+21 |
| FLN5 | 718F_706Phe | -61.91 | -0.09 | 7.11 | 0.29 | 45.77 | 6.6 | -2.49E+21 | 7.20E+20 |
| FLN5 | 726F_746Phe | -62.52 | -0.7 | 5 | 0 | 4.77 | 0.46 | -1.12E+22 | 1.05E+20 |
| FLN5 | 728F_744His | -62.39 | -0.57 | 5.38 | 0.06 | 20.9 | 0.47 | -7.40E+21 | 1.51E+20 |
| FLN5 | 728F_744Phe | -61.79 | 0.03 | 5.16 | 0.03 | 20.67 | 0.71 | -7.06E+21 | 2.41E+20 |
| FLN5 | 732F_692Trp | -61.7 | 0.12 | 7.49 | 0.19 | 71.23 | 5.58 | 3.82E+20 | 2.41E+20 |
| FLN5 | 732F_692Phe | -61.69 | 0.13 | 6.21 | 0.36 | 40.77 | 7.44 | -9.73E+20 | 3.28E+20 |
| FLN5 | 732F_692His | -61.62 | 0.2 | 5.6 | 0 | 32.7 | 0.69 | -2.05E+21 | 1.87E+19 |
| FLN5 | 740F | -62.06 | -0.24 | 9.52 | 0.07 | 10.2 | 1.01 | -1.26E+21 | 3.29E+19 |
| FLN5 | 748F | -61.85 | -0.03 | 9.55 | 0.02 | 21.87 | 3.49 | -6.27E+20 | 6.15E+19 |
| FLN4 | 555F | -62.81 | -0.99 | 5.7 | 0.46 | 13.4 | 5.45 | -1.35E+22 | 1.20E+21 |
| FLN4 | 565F_646His | -62.05 | -0.23 | 5.28 | 0.01 | 10.27 | 0.32 | -8.15E+21 | 1.03E+20 |
| FLN4 | 616F_571His | -62.08 | -0.26 | 5.4 | 0.14 | 8.53 | 1.53 | -8.59E+21 | 4.12E+20 |
| FLN4 | 624F_644Phe | -62.52 | -0.7 | 5.14 | 0.01 | 2.53 | 0.26 | -1.02E+22 | 1.64E+20 |
| I27 | 5F_24Phe | -61.71 | 0.11 | 8.54 | 0.27 | 31.37 | 9.53 | -1.29E+21 | 3.30E+20 |
| I27 | 6F_24Phe | -61.84 | -0.02 | 7.84 | 0.08 | 21.2 | 5.75 | -2.64E+21 | 3.66E+20 |
| I27 | 9F_22Phe | -61.66 | 0.16 | 10.26 | 0.77 | 68.37 | 1.38 | 6.76E+20 | 2.18E+20 |
| I27 | 14F_87Phe | -62 | -0.18 | 6.22 | 0.02 | 16.53 | 0.56 | -3.75E+21 | 1.20E+20 |
| I27 | 14F_87His | -62.15 | -0.33 | 6.07 | 0.02 | 3.2 | 1 | -6.95E+21 | 4.94E+19 |
| I27 | 20F_61Trp | -61.73 | 0.09 | 6.88 | 0.43 | 68.27 | 6.81 | 4.16E+20 | 1.81E+20 |
| I27 | 59F | -62.18 | -0.36 | 5.34 | 0.07 | 17.1 | 1.68 | -6.93E+21 | 2.73E+20 |
| I27 | 72F_35His | -62.17 | -0.35 | 5.86 | 0.35 | 22.5 | 8.75 | -5.13E+21 | 8.25E+20 |
| FLNa21 | 2242F | -62.2 | -0.38 | 6.58 | 0.58 | 15.53 | 5.9 | -5.51E+21 | 8.72E+20 |
| FLNa21 | 2244F | -61.11 | 0.71 | 6.27 | 0.35 | 44.6 | 13.84 | -2.69E+21 | 2.00E+21 |
| FLNa21 | 2258F_2296Phe | -62.1 | -0.28 | 5.63 | 0.11 | 21.43 | 1.74 | -8.08E+21 | 4.13E+20 |
| FLNa21 | 2306F_2322His | -62.53 | -0.71 | 4.78 | 0.01 | 4.07 | 0.4 | -1.16E+22 | 2.22E+20 |
| FLNa21-Mig | 2274F | -61.55 | 0.03 | 7.4 | 0.06 | 63.4 | 2.29 | 5.65E+20 | 9.10E+19 |
| FLNa21-Mig | 2274F_12His | -62.07 | -0.49 | 5.09 | 0.02 | 12.5 | 0.69 | -7.58E+21 | 4.67E+19 |

**Table S6**. $^{19}$F NMR chemical shifts and descriptors calculated from MD (C36m) predictions. Geometric factors (Geom.) are given in cm$^{-3}$. HemK and HRAS variants were not predicted with C36m due to protein instability observed for these wild-type proteins with C36m.

| Full dataset | | | | |
|---|---|---|---|---|
| | ColabFold | AF3 | ff15ipq | Combined |
| Total variants | 49 | 50 | 50 | 50 |
| Predicted positives | 20 | 22 | 21 | 31 |
| Predicted negatives | 29 | 28 | 29 | 19 |
| Actual positives | 29 | 30 | 30 | 30 |
| Actual negatives | 20 | 20 | 20 | 20 |
| TN | 16 | 18 | 19 | 25 |
| TP | 16 | 17 | 18 | 14 |
| FP | 4 | 4 | 2 | 6 |
| FN | 13 | 11 | 11 | 5 |
| TPR | 0.55 | 0.62 | 0.63 | 0.83 |
| TNR | 0.8 | 0.81 | 0.9 | 0.7 |
| FPR | 0.2 | 0.19 | 0.1 | 0.3 |
| FNR | 0.45 | 0.38 | 0.37 | 0.17 |
| PPV | 0.8 | 0.82 | 0.9 | 0.81 |
| NPV | 0.55 | 0.61 | 0.62 | 0.74 |
| FDR | 0.2 | 0.18 | 0.1 | 0.19 |
| FOR | 0.45 | 0.39 | 0.38 | 0.26 |

| C36m dataset | | | | |
|---|---|---|---|---|
| | ColabFold | AF3 | ff15ipq | C36m |
| Total variants | 42 | 42 | 42 | 42 |
| Predicted positives | 14 | 15 | 17 | 15 |
| Predicted negatives | 28 | 27 | 25 | 18 |
| Actual positives | 23 | 23 | 23 | 23 |
| Actual negatives | 19 | 19 | 19 | 19 |
| TN | 11 | 13 | 15 | 14 |
| TP | 16 | 17 | 17 | 18 |
| FP | 3 | 2 | 2 | 1 |
| FN | 12 | 10 | 8 | 9 |
| TPR | 0.48 | 0.57 | 0.65 | 0.61 |
| TNR | 0.84 | 0.89 | 0.89 | 0.95 |
| FPR | 0.16 | 0.11 | 0.11 | 0.05 |
| FNR | 0.52 | 0.43 | 0.35 | 0.39 |
| PPV | 0.79 | 0.87 | 0.88 | 0.93 |
| NPV | 0.57 | 0.63 | 0.68 | 0.67 |
| FDR | 0.21 | 0.13 | 0.12 | 0.07 |
| FOR | 0.43 | 0.37 | 0.32 | 0.33 |

**Table S7**. Performance of predictive $^{19}$F NMR ring current design strategy. TP = true positive; TN = true negative; FP = false positive; FN = false negative; TPR = true positive rate; FPR = false positive rate; TNR = true negative rate; FNR = false negative rate; PPV = positive predictive value; NPV = negative predictive value; FDR = false discovery rate; FOR = false omission rate. Positive and negative classifications and predictions are defined in the Methods section. The combined approach defines a positive prediction being predicted as a positive by at least one of the methods (ColabFold, AF3, ff15ipq MD) and a negative when predicted negative by all three methods.

| | | ColabFold | AF3 |
|---|---|---|---|
| True predictions | **Mean** | 97.3 | 87.7 |
| | **S.D.** | 2.5 | 11.3 |
| | **Min.** | 86.5 | 54.2 |
| | **Max.** | 98.8 | 97.9 |
| | **pLDDT > 70 (%)** | 100.0 | 88.9 |
| | **pLDDT > 90 (%)** | 96.9 | 61.1 |
| False predictions | **Mean** | 93.9 | 74.2 |
| | **S.D.** | 7.7 | 15.7 |
| | **Min.** | 66.3 | 52.0 |
| | **Max.** | 98.8 | 95.2 |
| | **pLDDT > 70 (%)** | 94.1 | 57.1 |
| | **pLDDT > 90 (%)** | 82.4 | 21.4 |

**Table S8**. Minimum pLDDT of labelling pairs (tfmF/tyrosine and second aromatic) for true and false predictions. pLDDT values were averaged over all five predicted models.
